# A neurotensin receptor type 1-derived pepducin acts as a biased allosteric modulator to regulate target receptor function

**DOI:** 10.1101/2025.10.14.682336

**Authors:** Rebecca L. Brouillette, Frédérique Lussier, Émile Breault, Nathan Meneboo, Magali Chartier, Élora Midavaine, Jérôme Côté, Véronique Blais, Christine E. Mona, Jean-Michel Longpré, Michel Grandbois, Martin Audet, Élie Besserer-Offroy, Philippe Sarret

## Abstract

Pepducins are synthetic membrane-tethered lipopeptides designed to allosterically modulate G protein-coupled receptor (GPCR) signaling. Here, we characterize a series of pepducins targeting the neurotensin receptor type 1 (NTS1), revealing multifaceted modulation of this receptor class. Using BRET-based biosensors, we show that PP-001, a pepducin derived from NTS1’s first intracellular loop, preferentially activates G protein over β-arrestin signaling while inhibiting NT binding, NT-induced β-arrestin recruitment, and NTS1 receptor internalization, thereby acting as biased allosteric agonist and negative allosteric modulator. PP-001 also promotes the formation of both homo- and heteromeric multi-receptor units. In vivo, PP-001 elicits potent, sustained hypotensive effects, reversible by the NTS1 antagonist SR48692. Finally, although the mechanism of pepducin-receptor interaction remains unclear, this study identifies a critical N-terminal RKK motif for PP-001’s biological activity. Thermodenaturation assays with purified NTS1 and mutagenesis further provide evidence for the role of NTS1’s H8 domain in direct pepducin-receptor interaction. This work highlights pepducins’ modulatory potential as pharmacological tools for GPCR-targeted drug development.

## 1. Introduction

Pepducins represent a unique class of drugs designed to target G protein-coupled receptors (GPCRs), also known as 7-transmembrane domain receptors (7TMRs), by using a distinctive intracellular allosteric mechanism. Structurally, pepducins consist of a lipid moiety, most commonly palmitate, conjugated to a peptide derived from one of the four intracellular domains (ICLs 1–3, C-terminus) of a receptor of interest ^1^. Functionally, the lipid serves as a tether to the cell membrane’s phospholipid bilayer and allows the pepducin’s passive translocation between the outer and inner leaflets ^2,3^. Once pepducin penetrates the intracellular compartment, it is thought to allosterically bind to its cognate receptor and thereby modulate the signaling output ^4^. However, the nature of this modulation may be multifarious. Some pepducins have been reported to directly promote 7TMR signaling and could thus be classified as allosteric agonists ^5–7^. Others have been shown to inhibit the activity of an extracellular agonist, behaving as negative allosteric modulators (NAMs) of their target receptor ^8–10^. Conceivably, they could also potentiate the activity of an extracellular agonist and behave as positive allosteric modulators (PAMs) of their receptor, although no examples of pepducins acting in this way have been reported. A small number of pepducins are also known to inhibit responses triggered by other (agonistic) pepducins ^11,12^. Importantly, these lipopeptides can display signaling bias by preferentially activating (or inhibiting) a subset of the receptor’s associated signaling pathways, unlike a balanced reference ligand ^6,7,13^. Finally, due to their unique mode of action, pepducins may regulate receptor function in ways that are simply not possible with traditional ligands ^14^. In short, they have the potential to be very useful tools in drug discovery, for both research and therapeutic purposes.

To date, pepducins have been derived from a range of 7TMRs, including protease-activated receptors 1, 2, and 4 (PAR1, PAR2, and PAR4) ^1,15^; chemokine receptors CCR2 ^16^, CXCR1/2 ^17,18^, and CXCR4 ^5,17^; β2 adrenergic receptor (β2R) ^6,7^; formyl peptide receptors 1, 2, and 3 [FPR1 ^19^, FPR2 ^20,21^, and FPR3 ^22^]; sphingosine-1-phosphate receptors S1PR1 and S1PR3 ^12,23^; and urotensin receptor (UTR) ^24^. With such a varied list of targets, the pathophysiological contexts in which pepducins have been studied are equally diverse: arthritis ^25^, cancer [breast ^26^, intestinal ^27,28^, lung ^8,29^, ovarian ^18,30^, prostate ^31^, and leukemia ^32^], heart failure ^33^, dermatitis and itch ^34,35^, diabetes ^36^, liver disease ^37^, sepsis ^17,38^, and thrombosis ^39,40^, to name a few. Among them, PZ-128, a PAR1-derived pepducin, has successfully completed Phase I ^41^ and Phase II ^39^ clinical trials, and may thus become a first-of-its class drug on the market. Despite these advances, many questions remain about pepducins, including the precise nature of the pepducin-receptor interaction, which is still unclear. Thus, there is a need to deepen our mechanistic understanding of how these lipopeptides operate and to broaden our set of available tools by applying the pepducin strategy to new targets. To this end, we recently introduced a new series of pepducins derived from the ICL1 of the neurotensin receptor type 1 (NTS1), one of the two 7TMRs bound by the neurotensin tridecapeptide (NT) ^42^. This Class A GPCR is expressed in central ^43^ and peripheral ^44,45^ tissues, and while its activation has been linked to a range of effects such as hypotension ^46^ and hypothermia ^47^, it is of particular interest for pain relief, as NT is known to produce potent opioid-independent analgesia when administered to rodents ^48–50^. Prior to our work, NTS1 had not been targeted by pepducin technology, and research into the therapeutic potential of pepducins in chronic pain paradigms was limited. Interestingly, we recently showed that the pepducin PP-001 derived from ICL1 exerted analgesic effects in various pain models ^42^. In the present study, we aim to characterize in depth the signaling profile of PP-001 at the NTS1 level and to investigate the molecular basis of pepducin-receptor interaction. Our results reveal complex modulation of NTS1 signaling, as PP-001 was found to behave as a biased allosteric agonist by favoring G protein signaling over βarrestin recruitment and as a negative allosteric modulator by decreasing NT binding and signaling. Our alanine scanning experiments on PP-001 and NTS1’s ICL domains also provide important insights into the molecular determinants governing the pepducin-receptor interaction. In particular, we identify that the N-terminal RKK motif of PP-001 plays a critical role in its biological activity and show that the eighth α-helix (H8) of NTS1 represents an important site at the interface of pepducin-receptor interaction. Finally, thermodenaturation assays with purified NTS1 receptor constructs demonstrated that PP-001 interacts directly with NTS1.

## 2. Materials and Methods

### 2.1. Materials

To perform the experiments described here, we synthesized the compounds NT, NT(8-13), and our NTS1-derived pepducins at the peptide synthesis facility of the Université de Sherbrooke (http://www.usherbrooke.ca/ips/en/platforms/psp/). These include the full-length ICL1 pepducin (NTS1-ICL1-PP-001), a non-palmitoylated control (NTS1-ICL1-NP-001), a pepducin with a scrambled peptide sequence (NTS1-ICL1-PP-SCR-001), a C-terminally truncated pepducin (NTS1-ICL1-PP-005), and a 13-pepducin alanine scan series in which the amino acids at positions 2 to 14 of NTS1-ICL1-PP-001 (corresponding to residues R90 to H102 of hNTS1) have been sequentially replaced by alanine residues (NTS1-ICL1-[Ala^2^]PP-001 to NTS1-ICL1-[Ala^14^]PP-001). We also obtained the NTS1 antagonist SR48692 from R&D Systems, Inc. (Minneapolis, MN, USA). The specific peptide sequences are provided in **Table 1**.

**Table 1:** Peptide sequences of tested compounds.

| Compound | Abbreviation | Sequence |
| --- | --- | --- |
| Neurotensin | NT | H<sub>2</sub>N– E L Y E N K P R R P Y I L –COOH |
| Neurotensin (8-13) | NT(8-13) | H<sub>2</sub>N– R R P Y I L –COOH |
| NTS1-ICL1-PP-001 | PP-001 | Palmitoyl– A R K K S L Q S L Q S T V H –CO-NH<sub>2</sub> |
| NTS1-ICL1-NP-001 | NP-001 | H<sub>2</sub>N– A R K K S L Q S L Q S T V H –CO-NH<sub>2</sub> |
| NTS1-ICL1-PP-SCR-001 | PP-SCR-001 | Palmitoyl– A R K K S L Q S L Q S T V H –CO-NH<sub>2</sub> |
| NTS1-ICL1-PP-005 | PP-005 | Palmitoyl– A R K K S L –CO-NH<sub>2</sub> |
| NTS1-ICL1-[Ala<sup>2</sup>]PP-001 | [Ala<sup>2</sup>]PP-001 | Palmitoyl– A **A** K K S L Q S L Q S T V H –CO-NH<sub>2</sub> |
| NTS1-ICL1-[Ala<sup>3</sup>]PP-001 | [Ala<sup>3</sup>]PP-001 | Palmitoyl– A R **A** K S L Q S L Q S T V H –CO-NH<sub>2</sub> |
| NTS1-ICL1-[Ala<sup>4</sup>]PP-001 | [Ala<sup>4</sup>]PP-001 | Palmitoyl– A R K **A** S L Q S L Q S T V H –CO-NH<sub>2</sub> |
| NTS1-ICL1-[Ala<sup>5</sup>]PP-001 | [Ala<sup>5</sup>]PP-001 | Palmitoyl– A R K K **A** L Q S L Q S T V H –CO-NH<sub>2</sub> |
| NTS1-ICL1-[Ala<sup>6</sup>]PP-001 | [Ala<sup>6</sup>]PP-001 | Palmitoyl– A R K K S **A** Q S L Q S T V H –CO-NH<sub>2</sub> |
| NTS1-ICL1-[Ala<sup>7</sup>]PP-001 | [Ala<sup>7</sup>]PP-001 | Palmitoyl– A R K K S L **A** S L Q S T V H –CO-NH<sub>2</sub> |
| NTS1-ICL1-[Ala<sup>8</sup>]PP-001 | [Ala<sup>8</sup>]PP-001 | Palmitoyl– A R K K S L Q **A** L Q S T V H –CO-NH<sub>2</sub> |
| NTS1-ICL1-[Ala<sup>9</sup>]PP-001 | [Ala<sup>9</sup>]PP-001 | Palmitoyl– A R K K S L Q S **A** Q S T V H –CO-NH<sub>2</sub> |
| NTS1-ICL1-[Ala<sup>10</sup>]PP-001 | [Ala<sup>10</sup>]PP-001 | Palmitoyl– A R K K S L Q S L **A** S T V H –CO-NH<sub>2</sub> |
| NTS1-ICL1-[Ala<sup>11</sup>]PP-001 | [Ala<sup>11</sup>]PP-001 | Palmitoyl– A R K K S L Q S L Q **A** T V H –CO-NH<sub>2</sub> |
| NTS1-ICL1-[Ala<sup>12</sup>]PP-001 | [Ala<sup>12</sup>]PP-001 | Palmitoyl– A R K K S L Q S L Q S **A** V H –CO-NH<sub>2</sub> |
| NTS1-ICL1-[Ala<sup>13</sup>]PP-001 | [Ala<sup>13</sup>]PP-001 | Palmitoyl– A R K K S L Q S L Q S T **A** H –CO-NH<sub>2</sub> |
| NTS1-ICL1-[Ala<sup>14</sup>]PP-001 | [Ala<sup>14</sup>]PP-001 | Palmitoyl– A R K K S L Q S L Q S T V **A** –CO-NH<sub>2</sub> |

Throughout this article, pepducins are referred to by their abbreviated titles PP-001, NP-001, PP-SCR-001, etc. The protected amino acids and TentaGel R-RAM resins used in their synthesis were obtained from Chem-Impex International (Wood Dale, IL, USA). All other chemicals and reagents involved in the synthesis of peptides and pepducins were obtained from Sigma-Aldrich (Oakville, ON, Canada) or Fisher Scientific (Montreal, QC, Canada). Dulbecco’s Modified Eagle’s Medium (DMEM), Dulbecco’s Modified Eagle Medium: Nutrient Mixture F12 (DMEM-F12), Leibovitz (L-15) medium, HEPES (4-(2-hydroxyethyl)-1-piperazineethanesulfonic acid), penicillin-streptomycin, fetal bovine serum (FBS), gentamycin G418, phosphate-buffered solution (PBS), Hank’s Balanced Salt Solution (HBSS), and trypsin (0.25%) supplemented with 0.53 mM EDTA were all obtained from Wisent Bioproducts (St. Bruno, QC, Canada). Opti-MEM Reduced Serum Medium was acquired from Invitrogen (Burlington, ON, Canada). Salmon sperm DNA (SSD) and polyethyleneimine (PEI) were sourced from Sigma-Aldrich (Oakville, ON, Canada). Coelenterazine 400A (Deep Blue C) was provide by GoldBio (Gold Biotechnology, St-Louis, MO, USA). White opaque 96-well plates were purchased from BD Falcon (Corning, NY, USA). 96-well plates with gold-plated interdigitated finger electrodes for Electric Cell-substrate Impedance Sensing (ECIS) were purchased from Applied Biophysics (Troy, NY, USA).

### 2.2. Pepducin synthesis

NTS1-derived pepducins were synthesized using standard C→N solid-phase peptide synthesis, with 12 mL polypropylene cartridges with 20 μm PE frit (Applied Separations, Allentown, PA, USA). 250 µmol of TentaGel S RAM resin (0.24 mmol/g) were deprotected with a mixture of piperidine:DMF (1:1) for 2 × 10 min. The fmoc-protected amino acids (3 eq.) were coupled using HATU (3 eq.) and DIEA (6 eq.) in DMF (10 mL) for at least 2 h. The reactant and solvent were then filtered, and the resin was washed with 10 mL DMF (2 × 5 min under agitation) and three alternating cycles of washing with 2-propanol (7 mL) or DCM (7 mL). Deprotection cycles were carried out with a mixture of piperidine: DMF (1:1) for 2 × 10 min. After deprotection, the solvent was removed by filtration, the resin was washed with DMF, 2-propanol, and DCM (as described above), and the Fmoc-protected amino acid was then coupled using HATU/DIEA in DMF. Coupling of palmitate (3 eq.) was done in dry N-Methyl-2-Pyrrolidone (NMP) with HATU (3 eq.) and DIEA (6 eq.).

Peptides and pepducins were cleaved from the resin using 92.5% TFA, 2.5% TIPS, 2.5% H2O, and 2.5% EDT for at least 2 hours to provide the crude mixture of the desired peptide. The resin suspension was filtered through cotton wool and the peptide precipitated in 50 mL of tert-butyl methyl ether at 0°C. The suspension was centrifuged for 20 min at 1500 rpm, and the pellet was resuspended in a mixture of water:acetonitrile (2:1) and lyophilized before purification.

Purification was done on a Waters Mass-triggered preparative HPLC system (Sample Manager 2767, Binary Gradient Module 2545, with two 515 HPLC pumps, a System Fluidics Organizer, and a 2998 Photodiode Array Detector) combined to a SQ Detector 2 and equipped with an X Select CSH Prep C_18_ 5 mm OBD 19 mm × 250 mm column using a 25–40% gradient of acetonitrile with 0.1% formic acid for 15 min. The fractions were analyzed by UPLC/MS (Water H Class Acquity UPLC, mounted with Acquity UPLC BEH C_18_ column, 1.7 μm, 2.1 mm × 50 mm, and paired to a SQ Detector 2) using a gradient of 5–95% acetonitrile with 0.1% formic acid for 2 min. Fractions with a purity of 95% or greater were then lyophilized and stored at -20°C until use. HRMS was performed on a Bruker MaXis 3G high-resolution Q-ToF. Purified product yielded a white powder after being freeze-dried. Stock solutions in 100% DMSO were prepared for biological assays at 10-mM concentrations.

### 2.3. Plasmids and constructs

The cDNAs encoding hNTS1 and the Gβ_1_ subunit were purchased from the Bloomsburg University cDNA Resource Center (Bloomsburg, PA, USA). Those encoding Gα_q_-RlucII (^51^), Gα_oA_-RlucII (^52^), Gα_13_-RlucII (^53^), GFP10-Gγ_1_ (^54,55^), RlucII-β-arrestin-1 (^56^) or -2 (^57^), and rGFP-CAAX (^58^) were kindly provided by Dr. Michel Bouvier (Université de Montréal). The plasmids encoding NTS1-GFP10 (^59^) and APJ-GFP10 (^60^) were previously constructed by our team using an InFusion advantage PCR cloning kit (Clontech Laboratories, Mountain View, CA, USA). To generate the NTS1-RlucII and APJ-RlucII constructs for this study, the complete coding sequences of hNTS1 and hAPJ receptors were amplified by PCR without their stop codons, inserted in frame into a pIRES(puroR)-RlucII-StrepTagII vector following BamHI digestion, and fused to RlucII by a 7 amino acid GSGSAGT linker using the InFusion advantage PCR cloning kit according to the manufacturer’s recommendations. Finally, a 3xHA-hNTS1 pcDNA3.1 construct was purchased from the Bloomsburg University cDNA Resource Center (Bloomsburg, PA, USA) and provided to GenScript (Piscataway, NJ, USA) as the template for the construction of the 27 NTS1 ICL mutant cDNAs (ICL Mutants 1–27). The specific mutations introduced are provided in **Supplementary Table S6**. Constructs were verified by DNA sequencing before use. Engineered recombinant NTS1-3 and NTS-22 were constructed using standard cloning procedures and polymerase chain reaction. The receptor modifications are presented in **Supplementary Fig. S2** and **Supplementary Table S5**, as described elsewhere (^61,62^).

### 2.4. Cellular assays

#### 2.4.1. Cell culture and transfections

Multiple cell lines were used to characterize the in vitro behavior of our pepducin series. Chinese hamster ovary cells (CHO-K1, CCL-61 from ATCC, Manassas, VA) were cultured in DMEM-F12 medium supplemented with 20 mM HEPES, 10% FBS, and penicillin (100 U/mL)-streptomycin (100 μg/mL). CHO-K1 cells stably expressing hNTS1 (CHO-hNTS1, ES-690-C), purchased from Perkin Elmer (Montreal, QC, Canada), were cultured in the same conditions but further supplemented with 0.4 mg/mL of geneticin (G418). Human embryonic kidney 293 (HEK293) cells were cultured in DMEM supplemented with 20 mM HEPES, 10% FBS, and penicillin (100 U/mL)-streptomycin (100 μg/mL). All cell lines were maintained in a humidified atmosphere at 37°C and 5% CO_2_ and were used between passages 5 and 25.

For transient expression of recombinant proteins, 100-mm^2^ cell culture dishes were seeded with 2 x 10^6^ cells. Cells were transfected 24 h later with 12 μg cDNA; if the amount of cDNA required for the assay did not reach this total, transfection mixes were completed with SSD. Transfection complexes were formed by incubating plasmids for 25 min in Opti-MEM serum-free media with the transfection reagent polyethyleneimine (PEI) at a 3:1 PEI:cDNA ratio (^63^), then gently deposited onto cells.

#### 2.4.2. Bioluminescence Resonance Energy Transfer (BRET) assays

##### 2.4.2.1. G protein dissociation and βarr recruitment

To monitor G protein dissociation, CHO-K1 cells were transfected with either Gα_q_-RlucII, Gα_oA_-RlucII, or Gα_13_-RlucII, as well as GFP10-Gγ_1,_ Gβ_1_, and hNTS1. To monitor βarr recruitment, CHO-K1 cells were transfected with RlucII-β-arrestin 1 or RlucII-β-arrestin 2 and NTS1-GFP10. At 24 h post-transfection, the cells were detached with trypsin (0.25%) containing 0.53 mM EDTA and plated (50,000 cells/well) in white opaque 96-well plates. At 48 h post-transfection, the cells were washed once with HBSS, and 90 μL of HBSS containing 20 mM HEPES was added (or 80 μL for the allosteric modulator assays). Cells were equilibrated for 1 h at 37°C. In the agonist assays, cells were stimulated for 10 min with increasing concentrations of NT(8-13) (10^-6^ to 10^-12^ M) or NTS1-derived pepducins (10^-4^ to 10^-8^ M), followed by coelenterazine 400A (5 μM). In the allosteric modulator assays, cells were first stimulated with fixed concentrations of the pepducins (0, 1, 10, 32, 100 μM) for 10 min, followed by increasing concentrations of NT or NT(8-13) (10^-6^ M to 10^-12^ M) and coelenterazine 400A (5 μM). Plates were read multiple times over a 20-min period, using a Mithras^2^ LB 943 plate reader (Berthold Technologies, Oak Ridge, TN, USA) with a filter set selected for BRET^2^ measurement (λem_RlucII_ 410 ± 80 nm; λem_GFP10_ 515 ± 40 nm) set to a 1-s integration time. The BRET^2^ ratio was set at 515:410 nm (GFP10 emission:RlucII-coelenterazine 400A emission).

##### 2.4.2.2. Receptor endocytosis

To monitor NTS1 receptor endocytosis, CHO-K1 cells were transfected with NTS1-RlucII and rGFP-CAAX using the transfection reagent PEI, then transferred into 96-well opaque white plates 24 h later, as described above. At 48 h post-transfection, cells were washed with HBSS, received 90 μL of HBSS containing 20 mM HEPES (or 80 μL for the allosteric modulator assays), and were equilibrated for 1 h at 37°C. In the agonist assays, cells were first stimulated with coelenterazine 400A (5 μM), and BRET^2^ luminescence was recorded on a Mithras^2^ LB 943 plate reader (Berthold Technologies, Oak Ridge, TN, USA) for 5 min at 20-s intervals, to acquire a baseline pre-pepducin stimulation. Cells were then stimulated with vehicle (HBSS, 1% DMSO), NT(8-13) (1 μM), or NTS1-derived pepducins (50 μM), and BRET^2^ was recorded for a further 30 min at 20-s intervals (0.5-s integration time) at 37^ο^C. In the allosteric modulator assays, cells were first stimulated with vehicle (HBSS, 1% DMSO) or with NTS1-derived pepducins (50 μM or 100 μM) and incubated at 37°C for 10 min, before receiving 5 μM of coelenterazine 400A. As above, BRET^2^ was recorded for 5 min at 20-s intervals before stimulation with HBSS or NT(8-13) (1 μM), then monitored for a further 30 min. BRET^2^ ratios for each condition were normalized to reflect a BRET^2^ change factor: BRET^2^ ratios at each time-point were compared to (divided by) baseline BRET^2^ ratios. These normalized responses were then compared to (subtracted from) those of the vehicle condition (or vehicle + HBSS condition).

##### 2.4.2.3. Receptor homo- and heteromerization

To monitor NTS1-NTS1 homomeric or NTS1-APJ heteromeric interactions, BRET^2^ titration experiments were performed using the biosensor couples hNTS1-RlucII and hNTS1-GFP10; hNTS1-RlucII and hAPJ-GFP10; or hAPJ-RlucII and hNTS1-GFP10. CHO-K1 cells (NTS1-NTS1 homomer experiments) or HEK293 cells (NTS1-APJ heteromer experiments) were seeded (300,000 cells/well) onto 6-well cell culture dishes and transfected 24 h later with transfection complexes. Each condition received a total of 2.1 μg cDNA, corresponding to fixed amounts (50, 100 ng) of 7TMR-RlucII and increasing amounts (0–2 μg) of 7TMR-GFP10. Transfection mixes were completed with SSD. At 24 h post-transfection, cells were detached with trypsin (0.25%) supplemented with 0.53 mM EDTA and plated (100,000 cells/well) into white opaque 96-well plates. At 48 h post-transfection, cells were washed once with HBSS and received 90 μL of HBSS supplemented with 20 mM HEPES. Cells were then equilibrated for 1 h at 37°C, after which they were stimulated with buffer, with NT(8-13) or Apelin-13 (1 μM), or with NTS1-derived pepducins (50 or 100 μM) for 10 min, followed by coelenterazine 400A (5 μM). Plates were read multiple times over a 20-min period, using a Mithras^2^ LB 943 plate reader (Berthold Technologies, Oak Ridge, TN, USA) with a filter set selected for BRET^2^ measurement (λem_RlucII_ 410 ± 80 nm; λem_GFP10_ 515 ± 40 nm). The BRET^2^ ratio was determined as GFP10em/RlucIIem. Baseline BRET^2^ ratios (condition with no transfected 7TMR-GFP10) were subtracted from each condition’s BRET^2^ ratios to obtain ΔBRET^2^.

### 2.5. Radioligand binding

To determine the pepducins’ ability to affect orthosteric NT binding at the NTS1 receptor, competitive radioligand binding assays were performed. CHO-hNTS1 cells were cultured as described above; once 80% confluency was reached, cells were washed with PBS and frozen at -80°C. Frozen cells were thawed, scraped from the dish with 10 mM Tris buffer with 1 mM EDTA (pH 7.5) and centrifuged at 15,000 g for 5 min at 4°C. The pellet was resuspended in binding buffer (50 mM Tris-HCl [pH 7.5], 0.2% BSA). 15 μg of cell membranes expressing the hNTS1 receptor were incubated with 45 pM ^125^I-[Tyr3]-NT (2200 Ci/mmol, purchased from PerkinElmer, Billerica, MA, USA) in the presence of increasing concentrations of NT(8-13) (10^-12^ to 10^-6^ M), PP-001, NP-001, PP-SCR-001, [Ala^2^]PP-001, or [Ala^13^]PP-001 (10^-8^ to 10^-4^ M) for 60 min at room temperature.

In the binding experiments performed with HA-hNTS1 and the ICL mutants (Mutants 1–27, whose sequences are shown in **Supplementary Table S6**), HEK293 cells were transfected with 6 μg of cDNA, completed to 12 μg with SSD. At 24 h post-transfection, cells were washed with PBS and frozen at -80°C. Cells were scraped, centrifuged, and resuspended in 1 mL of binding buffer without BSA (50 mM Tris-HCl [pH 7.5]). Cell membranes were quantified using the Pierce BCA Protein Assay Kit (Thermo Fisher Scientific, Montreal, QC, Canada) according to the manufacturer’s instructions. Cell membrane suspensions were re-prepared in 6 μg/well of binding buffer (50 mM Tris-HCl [pH 7.5], 0.2% BSA). ICL mutants were subdivided into 3 groups relative to their cell surface expression data: Group 1 (full cell surface expression), including Mutants 3, 6, 7, 9, 10, 11, 12, 13, 15, 21, 22, 23, 24, 25, 26, and 27; Group 2 (partial cell surface expression), including Mutants 2, 5, 8, 14, 16, 17, and 20; and Group 3 (poor cell surface expression), including Mutants 1, 4, 18, and 19. Cell membranes were incubated with ^125^I-[Tyr^3^]-NT (2200 Ci/mmol) in binding buffer in the presence of increasing concentrations of NT(8-13) (10^-12^ M to 10^-6^ M) or PP-001 (10^-6^ M to 10^-4^ M) for 60 min at room temperature; Group 1 cell membranes were incubated with 90 pM ^125^I-[Tyr^3^]-NT, Group 2 cell membranes with 180 pM ^125^I-[Tyr^3^]-NT, and Group 3 cell membranes with 300 pM ^125^I-[Tyr^3^]-NT.

After incubation, the binding reaction mixtures were transferred to PEI-coated 96-well filter plates (glass fiber filters GF/B, Millipore, Billerica, MA, USA). Reaction was terminated by filtration, and plates were washed three times with 200 μL of ice-cold binding buffer. Glass filters were then counted using a γ-counter (2470 Wizard2, PerkinElmer, Mississauga, ON, Canada). Nonspecific binding was measured in the presence of 10^-5^ M unlabeled NT(8-13) and represented less than 5% of total binding. Data were normalized according to NT(8-13) to control for unwanted sources of variation.

### 2.6. Electric Cell-substrate Impedance Sensing (ECIS)

To monitor whole-cell integrative responses to pepducin treatment, CHO-hNTS1 cells were seeded (35,000 cells/well, 300 μL/well) onto cysteine-stabilized, Poly-L-Lysine-coated 96-well plates with interdigitated finger gold-plated electrodes (96W20idf) purchased from Applied Biophysics (Troy, NY, USA) and used as recommended by the manufacturer. Cells were cultured in serum-containing media for 24 h, then serum-starved for 16–18 h prior to the experiment. On the day of the experiment, the cells were washed with PBS and received 90 μL of L-15 modification medium. The biophysical parameters of the cell monolayer (i.e., electrical resistance to a 4000-Hz single frequency AC current) were recorded with an ECIS Zθ linked to a 96-well array station (Applied Biophysics, Troy, NY, USA) for a minimum period of 60 min prior to compound stimulation, to acquire a stable baseline. Cells were treated with L-15 media or 10 μM of PP-001, PP-SCR-001, or the PP-001 alanine scan series ([Ala^2^]PP-001, [Ala^3^]PP-001, [Ala^4^]PP-001, … [Ala^14^]PP-001). The dynamic mass redistribution (DMR) dynamics were then recorded for a further 60 min post-compound stimulation. The electrical resistivity traces shown represent the averaged normalized response, calculated as the resistance of the cell monolayer (ohms) divided by the average baseline resistance (ohms) prior to compound stimulation.

### 2.7. Cell surface expression of NTS1 ICL mutants

To determine the cell surface expression of the 27 NTS1 ICL mutants (whose sequences are shown in **Supplementary Table S6**), ELISA was conducted. HEK293 cells were seeded (100,000 cells/well) into 24-well plates coated with poly-L-Lysine and transfected with cDNA (415 ng/well) coding for WT HA-hNTS1 receptor or HA-hNTS1 ICL mutants (Mutants 1–27) using PEI (3:1 PEI:cDNA). Then, 48 h post-transfection, cells were fixed with 3.7% (v/v) formaldehyde in Tris-buffered saline (TBS, 20 mM Tris-HCl [pH 7.5], and 150 mM NaCl) for 5 min at room temperature. Cells were washed three times with TBS and incubated for 45 min in TBS supplemented with 3% (m/v) fat-free dry milk to block nonspecific binding sites. A mouse monoclonal anti-HA antibody coupled to HRP (Roche) was added at a 1:1000 dilution in TBS 3% fat-free milk and incubated for 2 h at room temperature on a nutating mixer. Following incubation, cells were washed three times with TBS before receiving 250 μL of 3,3’5,5’-Tetramethylbenzidine (TMB, Sigma-Aldrich, Oakville, ON, Canada). Plates were incubated for 15 min at room temperature, or until the colorimetric reaction (colorless to blue) was complete for the positive control (WT). The reaction was stopped by adding 250 μL of HCl 2N. At that point, 200 μL of the yellow reaction product was transferred to 96-well plates, and the absorbance was read at 450 nm on a Mithras^2^ LB 943 plate reader (Berthold Technologies, Oak Ridge, TN, USA). Cells transfected with the empty pcDNA3.1+ vector (mock) were used to determine background.

### 2.8. Expression and protein purification of NTS1-3 and NTS1-22

NTS1-3 and NTS1-22 were expressed in *Sf9* insect cells using Bac-To-Bac Bacculovirus Expression System (Thermo Fisher Scientific). Membrane form cells expressing NTS1-3 or NTS1-22 were prepared using three rounds of washing and ultracentrifugation at 250,000 g, first in the presence of lysis buffer containing 10 mM HEPES (pH 7.5), 20 mM KCl, and 10 mM MgCl_2_, and twice with a washing buffer containing 10 mM HEPES (pH 7.5), 1M NaCl, 20 mM KCl, and 10 mM MgCl_2_. The purified membrane was then resuspended in the lysis buffer supplemented with 20% (v/v) glycerol. 1 mM benzamidine and 5 µg/mL leupeptin were added for the first two steps. The membranes were then incubated for 1 h at 4°C with protease inhibitors, 1 mg/mL iodoacetamide, and 15 µM NT(8-13). The receptor was then solubilized in 50 mM HEPES (pH 7.5), 800 mM NaCl, 0.5% (w/v) n-dodecyl-β-D-maltopyranoside (DDM, Anatrace, Maumee, OH, USA), and 0.1% (w/v) cholesteryl hemisuccinate (CHS, Sigma-Aldrich) in the presence of protease inhibitors. The supernatant was isolated by ultracentrifugation for 1 h at 350,000 g and then incubated with TALON resin (TALON Superflow beads, Cytiva, Wilmington, DE, USA) overnight at 4°C. The TALON resin was washed with 20 column volumes of wash buffer 1 containing 50 mM HEPES (pH 7.5), 150 mM NaCl, 20 mM MgCl_2_, 20 mM imidazole, 8 mM ATP, 10% (v/v) glycerol, 0.1% (w/v) DDM, 0.02% (w/v) CHS, and 15 µM NT(8-13), followed by 10 column volumes of wash buffer 2 containing 50 mM HEPES (pH 7.5), 150 mM NaCl, 30 mM imidazole, 10% (v/v) glycerol, 0.05% (w/v) DDM, 0.01% (w/v) CHS, and 15 µM NT(8-13). The receptor was eluted using 2.5 column volumes of elution buffer containing 50 mM HEPES (pH 7.5), 150 mM NaCl, 250 mM imidazole, 10% (v/v) glycerol, 0.01% (w/v) DDM, 0.002% (w/v) CHS, and 15 µM NT(8-13). Finally, an Amicon Ultra 0.5 mL 100K (Millipore, Oakville, ON, Canada) was used to concentrate NTS1-3 and NTS1-22 at a concentration of 5 mg/ml.

### 2.9. Thermodenaturation assays

7-diethylamino-3-(4-maleimidophenyl)-4-methylcoumarin (CPM) was obtained from Thermo Fisher Scientific (Mississauga, ON, Canada) and resuspended in DMSO at a concentration of 500 µM. The assay used 30 or 40 μg of purified NTS1-3 or NTS1-22, respectively. The protein was added to a reaction buffer containing 50 mM HEPES (pH 7.5), 150 mM NaCl, 0.05% (w/v) DDM, 0.001% (w/v) CHS, and 10 µM CPM. The mixture was incubated in the presence of 100 µM of indicated ligand or vehicle for 20 min at room temperature. Unfolding of NTS1 was induced using a Rotor-Gene Q (Qiagen. Bay St., ON, Canada) thermocycler by slowly increasing the sample temperature (+0.5°C per step from 30°C to 95°C), resulting in an emission of a fluorescence signal detected in the blue channel (excitation 365 nm / emission 460 nm) from CPM chemical reaction to newly exposed receptor cysteines (^64^). The melting temperature was determined by the maximum of the first derivative of the normalized thermodenaturation curve using GraphPad Prism 10.

### 2.10. Blood pressure

The animal experimental procedures approved by the Ethical and Animal Care Committee of the Université de Sherbrooke (protocol 035-18B) were conducted in accordance with the policies and directives of the Canadian Council on Animal Care, and followed the ARRIVE Guidelines. Adult male Sprague-Dawley rats (250–300 g) were purchased from Charles River Laboratories (Saint-Constant, QC, Canada) and kept on a 12 h light / 12 h dark cycle with ad libitum access to food and water.

To monitor pepducin-induced changes in rat arterial blood pressure, male Sprague-Dawley rats were anesthetized with a mixture of ketamine and xylazine (87:13 mg/kg, i.m.) and placed in a supine position on a heated thermostatic pad. A PE50 catheter with heparinized saline was inserted into the rat’s right carotid artery and connected to a Micro-Med transducer (model TDX-300) linked to a blood pressure Micro-Med analyzer (model BPA-100c). Another PE10 catheter with heparinized saline was inserted into the left jugular vein for intravenous injection of compounds. Mean, systolic, and diastolic blood pressure, as well as heart rate, were recorded at 1-s or 5-s intervals for the duration of the test. Once basal blood pressure was stable, rats received 10-s bolus injections (1 mL/kg) of saline, vehicle (saline, 5% DMSO), or 55 nmol/kg dose solutions of NT(8-13), PP-001, NP-001, PP-SCR-001, [Ala^2^]PP-001, and [Ala^13^]PP-001. In the SR48692 + PP-001 group, rats were injected with the non-peptide antagonist SR48692 (1 mg/kg) 10 min prior to PP-001 administration. ΔMABP was determined relative to baseline. The minimum blood pressure relative to baseline (maximum ΔMABP) for each individual animal was determined, and values were pooled.

### 2.11. Statistical analysis

All data obtained for this study were plotted onto graphs using GraphPad Prism 10 software (RRID:SCR_002798) and represent the mean ± SEM of multiple independent experiments. Specific *n* values are provided in the figure legends. Sigmoidal concentration-response curves were plotted using the “log(agonist) vs. response (three-parameters)” regression or, in the case of the binding experiments, using the non-linear regression “One-site-Fit Log(IC_50_)”. pEC50 (or pIC_50_) and E_max_ values were compared using one-way ANOVA tests with Dunnett’s correction for multiple comparisons. Titration curves were plotted using the “one-site – specific binding” regression for binding saturation experiments. Similarly, BRET_50_ and BRET_max_ values were compared using one-way ANOVA with Dunnett’s correction. Otherwise, the details for data normalization, statistical analysis, and *P* values are provided in the figure legends.

## 3. Results

### 3.1. PP-001 preferentially promotes G protein activation over βarr recruitment

We recently designed and synthesized a pepducin series derived from the first intracellular loop (ICL1) of hNTS1 ^42^. This series included a full-length ICL1 pepducin (NTS1-ICL1-PP-001), a non-palmitoylated control (NTS1-ICL1-NP-001), and a pepducin with a scrambled peptide sequence (NTS1-ICL1-PP-SCR-001), which will be referred to throughout this paper by the abbreviations PP-001, NP-001, and PP-SCR-001, respectively. Their sequences, along with those of all peptides and lipopeptides tested throughout this study are provided in **Table 1**. Importantly, pepducin PP-001 has been shown to be effective in reversing nociceptive behaviors in rats, even in chronic neuropathic and inflammatory pain paradigms ^42^. Here, we have thoroughly investigated the signaling profile of PP-001 by evaluating the activation of Gαq, Gα_oA_, Gα_13_, βarr1, and βarr2 signaling pathways in NTS1-expressing cells in response to a range of pepducin concentrations.

To do so, we relied on bioluminescence resonance energy transfer (BRET^2^)-based cellular assays designed to monitor G protein dissociation and βarr recruitment, the principles of which are illustrated in **Fig. 1a** and **Fig. 1b**, respectively [Note that these schemas depict the classical mode of 7TMR activation by an extracellular, orthosteric ligand, whereas pepducins are thought to modulate their target receptor by acting on an intracellular allosteric site ^14^]. These assays, which monitor the interaction between donor-tagged (RlucII) and acceptor-tagged (GFP10) proteins, were performed 30 minutes after stimulation with pepducins in transiently transfected CHO-hNTS1 cells. The BRET^2^ ratios obtained were compared to NT’s C-terminal hexapeptide, NT(8-13). This hexapeptide represents the minimal biologically active fragment of NT ^65^ and has been shown to exhibit a signaling signature identical to that of NT in CHO-hNTS1 cells ^59^. NT(8-13) is therefore used as the reference compound in this study.

**Fig. 1.**
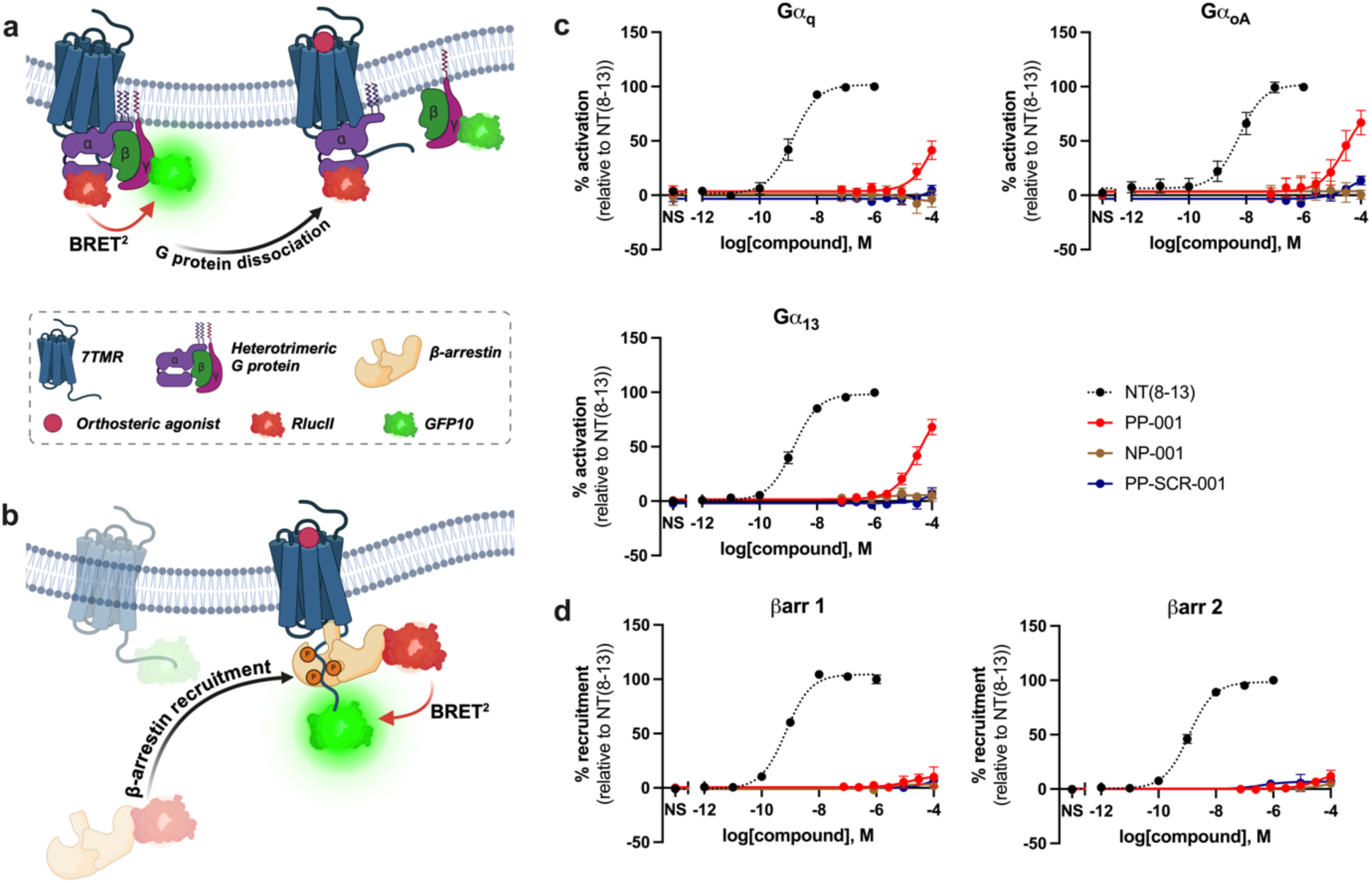
PP-001 activates G protein signaling pathways but does not recruit βarrs to NTS1. (a) Principle of the BRET^2^ G protein dissociation assay. Agonist-induced G protein activation leads to a loss of BRET^2^ signal, as donor-tagged (RlucII) Gα proteins and acceptor-tagged (GFP10) Gγ proteins dissociate. (b) Principle of the BRET^2^ βarr recruitment assay. The BRET^2^ signal increases as donor-tagged (RlucII) cytosolic βarrs are recruited to the acceptor-tagged (GFP10) receptor, after agonist binding. (c) Gα protein dissociation in CHO-K1 cells transiently expressing the NTS1 receptor, monitored by BRET^2^, in response to increasing concentrations of NT(8-13), PP-001, NP-001, or PP-SCR-001. (d) βarr recruitment in CHO-K1 cells, monitored by BRET^2^, in response to increasing concentrations of NT(8-13), PP-001, NP-001, or PP-SCR-001. Data represent the mean ± SEM of at least three independent experiments, tested in triplicate. Luminescence was measured 30 min after pepducin stimulation. BRET^2^ ratios were normalized to NT(8-13): data from vehicle-treated cells were set as 0% activation (or recruitment) and data from NT(8-13) (1 μM)-treated cells were set to 100% activation (or recruitment). Schemas were created with Biorender.com. NS: non-stimulated (treated with vehicle only).

As shown in **Fig. 1c** and **Fig. 1d**, our results are consistent with a biased signaling paradigm. Indeed, stimulation with PP-001 led to a marked increase in signaling pathway activation in the G protein dissociation assays. Maximum values of 41 ± 9% (Gα_q_), 67 ± 11% (Gα_oA_), and 68 ± 7% (Gα_13_) pathway activation were obtained at 100 μM PP-001. However, no recruitment of βarr1 and βarr2 at NTS1 was observed at the same concentration. Importantly, neither NP-001 nor PP-SCR-001 induced any change in BRET^2^, and thus do not appear to affect G protein signaling or βarr recruitment. Given that none of the G protein-related concentration-response curves could be fitted to a complete sigmoidal regression, including a second maximal plateau, we were unable to determine reliable potencies (EC_50_) for PP-001. Without these reliable potencies, we were also unable to do traditional bias quantifications using approaches based on determining intrinsic relative activity (RA_i_) values ^66,67^ or comparing transduction coefficients (τ/K_a_) ^68^ derived from the Black and Leff operational model ^69^. However, the complete lack of PP-001-mediated βarr recruitment at 100 μM strongly suggests that PP-001 behaves as a G protein-biased allosteric agonist.

### 3.2. PP-001 inhibits NT orthosteric binding and NT-induced βarr recruitment

PP-001 appears to act as a biased allosteric agonist of NTS1, preferentially activating G protein signaling pathways over βarr recruitment. However, it is now recognized that allosteric ligands can assume multiple identities, leading to classifications such as “ago-PAM” or “ago-NAM” (i.e., compounds that act as allosteric agonists in one signaling pathway, but as PAMs or NAMs in another) ^70^. As a result, we sought to determine whether treatment with PP-001 also affects NT binding at the orthosteric site or the activation of NT-promoted signaling pathways.

First, we tested PP-001 in a competitive radioligand binding assay. For this purpose, CHO-hNTS1 cell membrane preparations were incubated with radiolabeled NT ([^125^I]-NT) in the presence of increasing concentrations of unlabeled ligands. As shown in **Fig. 2a**, the pepducin PP-001 inhibited radiolabeled NT binding in a concentration-dependent manner, with an IC_50_ of 5.5 ± 0.8 μM and complete inhibition at 100 μM. As expected, NT(8-13) displaced radiolabeled NT with an IC_50_ of 1.0 ± 0.1 nM. Importantly, the peptide sequence of PP-001 alone (NP-001) had no effect on NT binding, suggesting that this “displacement” induced by PP-001 is not due to direct competition with NT at the orthosteric site, but results from the insertion of PP-001 into the membrane and its allosteric action. PP-SCR-001 was largely inactive, but slighly reduced NT binding at 100 μM.

**Fig. 2.**
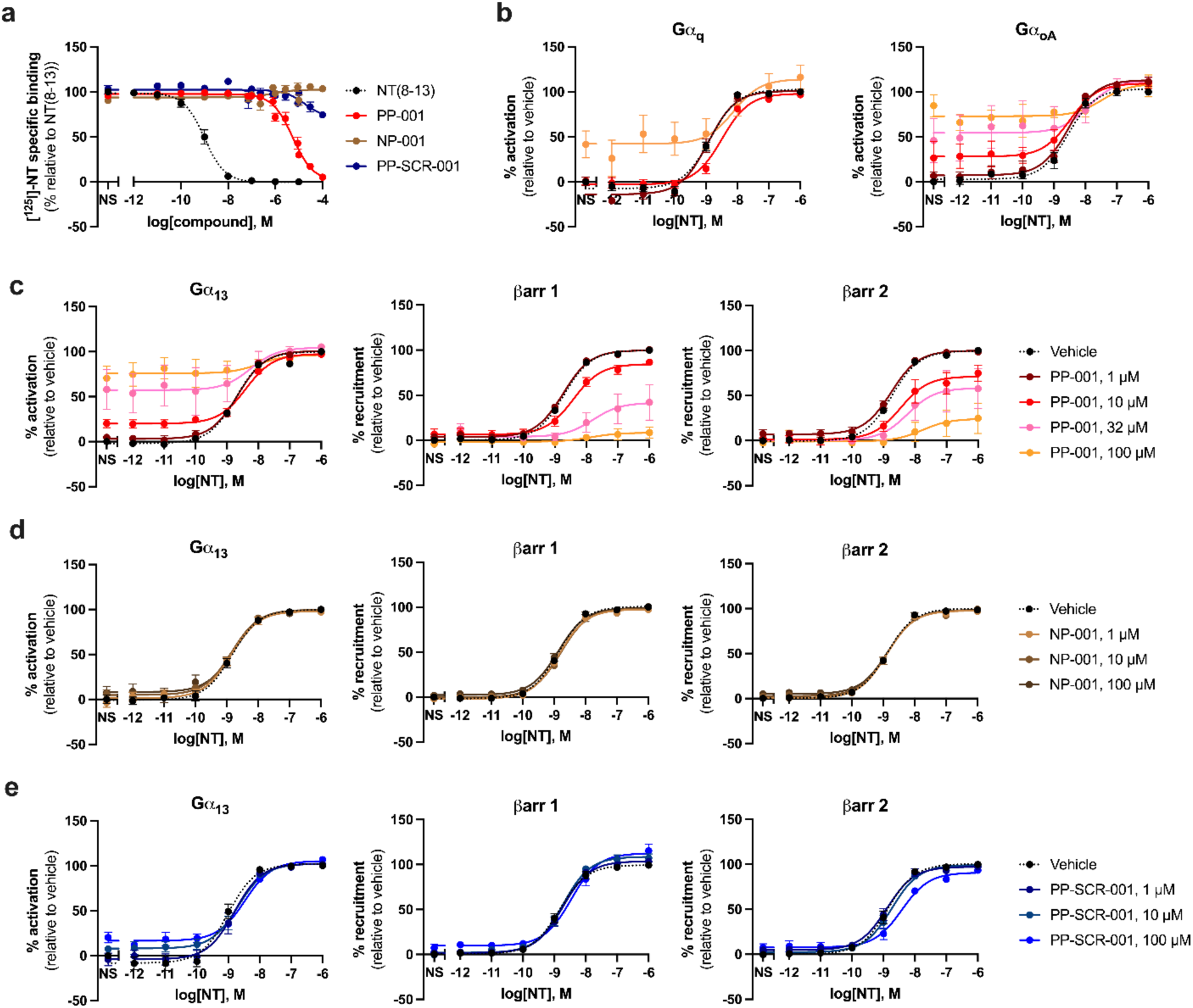
PP-001 allosterically inhibits NT binding and NT-promoted βarr recruitment, but not NT-promoted G protein activation. (a) Displacement of ^125^I-radiolabeled NT following incubation of CHO-hNTS1 membranes with increasing concentrations of unlabeled NT(8-13), PP-001, NP-001, or PP-SCR-001 in a competitive radioligand binding assay. (b, c) NT-induced G protein dissociation and βarr recruitment in CHO-K1 cells expressing NTS1, pre-treated (10 min) with increasing concentrations (0, 1, 10, 32, and 100 μM) of PP-001, as monitored by BRET^2^. (d-e) NT-induced Gα_13_ dissociation and βarr 1 and 2 recruitment in CHO-K1 cells pre-treated (10 min) with increasing concentrations (0, 1, 10, and 100 μM) of NP-001 and PP-SCR-001 (e), as monitored by BRET^2^. Luminescence was measured 30 min after pepducin stimulation, 20 min following NT(8-13) stimulation. Data represent the mean ± SEM of at least three independent experiments, tested in triplicate. Gamma radiation counts were normalized according to NT(8-13): data from untreated samples were set to 0% specific binding and data from NT(8-13) (1 μM)-treated samples were set to 100 % specific binding. BRET^2^ ratios were normalized according to NT(8-13), in the “vehicle” pre-stimulation condition: data from vehicle-treated cells were set to 0% activation (or recruitment) and data from cells treated with NT(8-13) (1 μM) were set to 100% activation (or recruitment). NS: non-stimulated (treated with buffer only).

Second, we performed BRET^2^ experiments to monitor G protein dissociation and βarr recruitment in an allosteric modulator-type assay, in which full concentration-response curves of NT were generated in cells pre-treated with fixed, but increasing concentrations of pepducins. As shown in **Fig. 2b** and **Fig. 2c**, pre-treating cells with PP-001 led to divergent effects in G protein versus βarr assays. In G protein dissociation assays, PP-001 increased the basal levels of pathway activation in a concentration-dependent manner, consistent with its previously determined allosteric agonist action. Treatment with PP-001 did not change NT’s E_max_ in a statistically significant manner (**Supplementary Table S1**). Additionally, although we observed a slight shift in NT’s EC_50_ values in each G protein pathway, this was accompanied by a corresponding increase in SEM. Therefore, we cannot conclude that NT’s potency was affected by PP-001 pre-treatment. If PP-001 inhibited NT-mediated signaling, as might be expected from the radioligand binding experiments, this result was effectively masked by its agonist action on G protein signaling.

In the βarr assays, however, there was a clear inhibitory effect on NT-mediated recruitment. A concentration-dependent decrease in the efficacy (E_max_) of NT to recruit both βarr1 and βarr2 was observed, reaching statistically significant levels at 10 μM of PP-001 and above. At 100 μM of PP-001, NT-mediated recruitment was almost completely abolished,with inhibition of 91 ± 7% and 80 ± 20% for βarr1 and βarr2, respectively (**Supplementary Table S1**). As with the G protein assays, we cannot conclude that the potency of NT to recruit βarrs was inhibited by PP-001, as the flattening of the NT curves led to a higher degree of uncertainty in the calculated EC_50_ values. Nevertheless, comparison of the pEC_50_ values between PP-001-treated and vehicle-treated conditions revealed a statistically significant rightward shift in NT potency, especially for βarr1 (**Supplementary Table S1**). Once again, the negative control pepducins NP-001 and PP-SCR-001 had no effect on the potency or efficacy of NT (**Fig. 2d** and **2e, Supplementary Table S2**); the pre-treated and vehicle-treated NT curves were superimposable in the G protein (Gα_13_) and βarr assays. As an additional control, we tested palmitate in the same experimental paradigm to ensure that a high concentration of lipids inserted into the cell membrane could not alone disrupt NTS1 signaling. NT-promoted signaling was not affected by palmitate pre-treatment (**Supplementary Table S2**). Overall, these results suggest that, in addition to acting as a biased allosteric agonist of NTS1, PP-001 acts as a NAM against the orthosteric NT binding and βarr signaling pathways.

### 3.3. PP-001 inhibits NT-induced receptor internalization

To support the results showing that PP-001 not only does not promote βarr recruitment to NTS1 but also inhibits NT-induced βarr recruitment, we sought to evaluate the effect of PP-001 on NTS1 receptor internalization. As it has been well established that βarr recruitment is a key step in the process of NTS1 endocytosis ^71–73^, we expected PP-001 treatment to mirror the results described above (i.e., not promote receptor internalization itself but inhibit NT-promoted receptor internalization). To monitor NTS1 receptor endocytosis, we used a BRET^2^ assay in which CHO-K1 cells were co-transfected with the donor-tagged NTS1 receptor (hNTS1-RlucII) and an acceptor-tagged plasma membrane marker (rGFP-CAAX) ^58^. The principle of the assay is schematized in **Fig. 3a**; an internalization of the receptor into the cell is reflected by a loss in BRET^2^ signal as the donor-tagged receptor becomes isolated from the membrane-bound acceptor sensor.

**Fig. 3.**
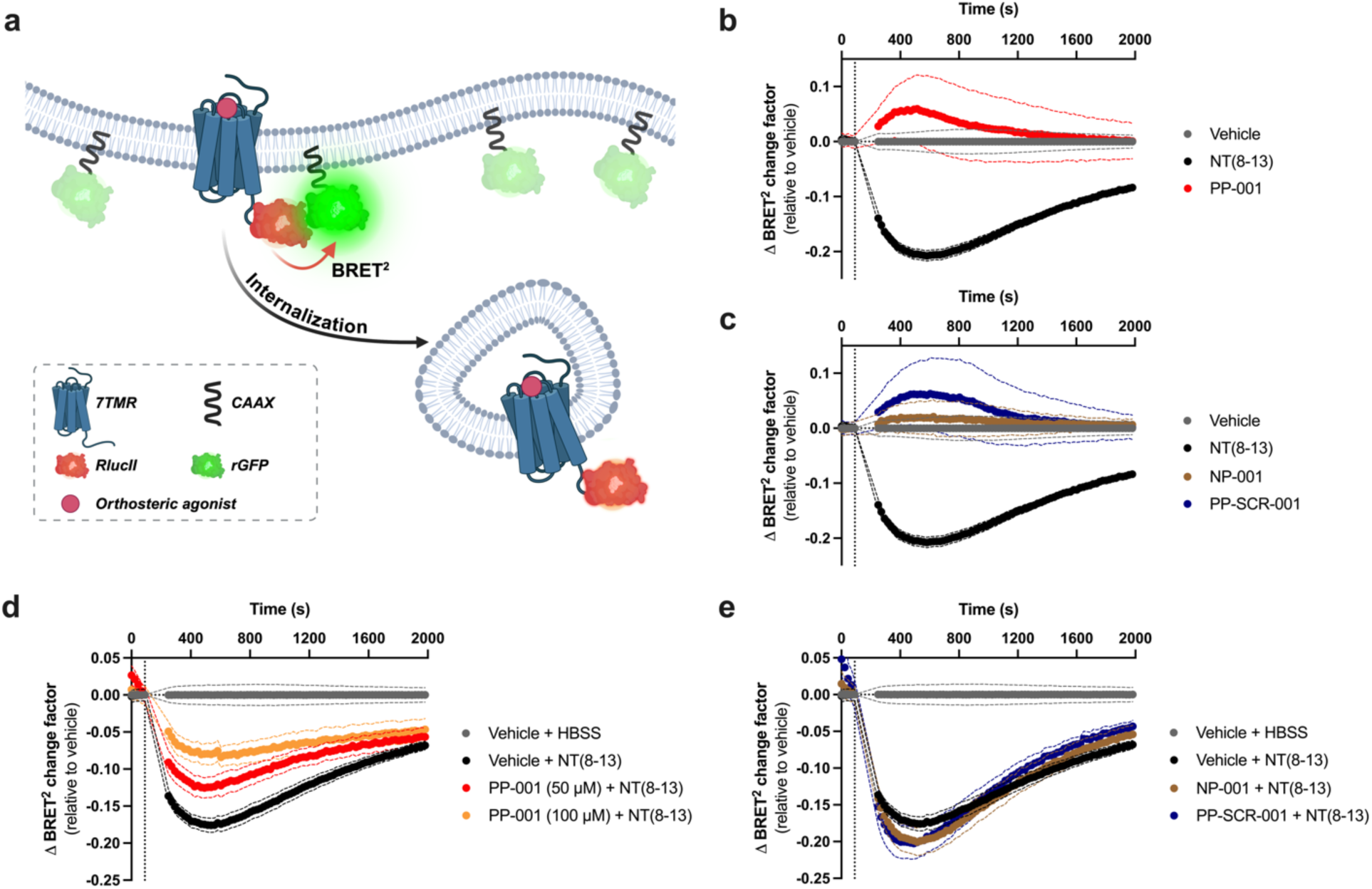
PP-001 does not promote NTS1 receptor internalization on its own but inhibits NT-induced receptor internalization. (a) Principle of receptor internalization assay. Agonist-induced receptor endocytosis leads to a loss of BRET^2^ signal, as the donor-tagged (RlucII) receptor is isolated from the acceptor-tagged, PM-localized CAAX-rGFP sensor. (b, c) Kinetic assays monitoring NTS1 receptor internalization over a 30 min period in transiently transfected CHO-K1 cells, following stimulation with vehicle (HBSS, 1% DMSO), NT(8-13) (1 μM), PP-001 (50 μM), NP-001 (50 μM) or PP-SCR-001 (50 μM). (d, e) Kinetic assay monitoring NT(8-13) (1 μM)-induced NTS1 receptor internalization over a 30 min period, in transiently transfected CHO-hNTS1 cells pre-stimulated (15 min) with PP-001 (50 μM, 100 μM), NP-001 (50 μM), or PP-SCR-001 (50 μM). Data represent mean ± SEM of four independent experiments, tested in duplicate. To determine a “ΔBRET^2^ change factor”, each condition was normalized according to itself and corrected to vehicle: baseline data for each condition was set as a value of 1 and subtracted from vehicle (or vehicle + HBSS)-treated condition. Schema was created with Biorender.com.

We first performed agonist type kinetic assays in which, following a baseline BRET^2^ luminescence reading, CHO-hNTS1 cells were stimulated with either the vehicle or a single concentration of NT(8-13) (1 μM), PP-001 (50 μM), NP-001 (50 μM), or PP-SCR-001 (50 μM). Luminescence was subsequently monitored over a 30-min period. As can be observed in **Fig. 3b** and **Fig. 3c**, NT(8-13) triggered a decrease in the BRET^2^ change factor, which is consistent with receptor endocytosis. A maximum decrease of 0.21 ± 0.01 BRET^2^ change factor was observed by 8.2 min post-treatment, followed by a slow return toward baseline. In contrast, PP-001 treatment did not produce any decrease. Rather, a slight increase was observed, although this was transient, highly variable between replicates, and also occurred in PP-SCR-001-treated cells. Compared to the vehicle, NP-001 did not trigger any change to the BRET^2^ ratio.

Subsequently, we performed allosteric modulator type kinetic assays, in which CHO-hNTS1 cells were pre-treated with the vehicle or pepducins for 10 min. As above, this step was followed by a baseline luminescence reading, stimulation with the vehicle or NT(8-13) (1 μM), and a 30-min readout period. As can be observed in **Fig. 3d**, pre-treatment of cells with PP-001 effectively reduced the NT(8-13)-promoted decrease in BRET^2^ ratio. Comparing the maximum decrease values in each condition, this corresponds to an inhibition of NT(8-13)-promoted NTS1 endocytosis of 43 ± 9% (50 μM) and 67 ± 7% (100 μM). The control compounds NP-001 and PP-SCR-001 did not reduce NT(8-13)-promoted endocytosis (**Fig. 3e**). Along with the βarr recruitment data, these findings reinforce the classification of PP-001 as a NAM, capable of inhibiting NT-mediated βarr recruitment and NTS1 receptor internalization in a concentration-dependent manner.

### 3.4. PP-001 may allosterically promote NTS1-NTS1 and NTS1-APJ interactions

To further characterize PP-001-mediated actions in vitro, we sought to evaluate whether PP-001 treatment impacts NTS1’s ability to self-associate into homomers or to form heteromers with other 7TMRs. Indeed, it has been established that 7TMRs can assemble to form dimers and higher-order oligomers, particularly for obligate dimer Class C GPCRs, although the relevance of this process in vivo is still debated ^74^. NTS1, specifically, has been shown to homomerize in detergent solutions ^75^, in reconstituted phospholipid bilayer systems ^76^, and in HELA cells ^77^. It has been proposed that this interaction is transient and follows a “rolling interface” model in which the protomers may adopt multiple configurations during their dimer lifetime ^78^.

To monitor NTS1-NTS1 interactions in live cells, we performed BRET^2^ titration experiments in which CHO-K1 cells were transfected with a fixed quantity of donor-tagged receptor (hNTS1-RlucII) and increasing quantities of acceptor-tagged receptor (hNTS1-GFP10) (**Fig. 4a**). In this experimental paradigm, specific receptor-receptor interactions are reflected by a BRET^2^ increase that follows a saturation curve, whereas nonspecific interactions are reflected by a linear BRET^2^ increase ^79^. As shown in **Fig. 4b** (left panel), a clear saturation curve was obtained under vehicle-treated condition, indicative of constitutive dimerization of NTS1 [conceivably, NTS1 may also assemble into higher-order complexes, although studies suggest that NTS1 predominantly forms dimers ^75,76,78^]. Treatment with pepducin PP-001 led to a significant upward shift, corresponding to a 25 ± 1% increase in BRET_max_ (100 μM, left panel of **Fig. 4b**; **Supplementary Table S3**). Interestingly, PP-001 did not affect the ratio of transfected GFP10/RlucII at which 50% BRET^2^ was observed (BRET_50_), suggesting that the pepducin may not increase the affinity of NTS1 protomers for each other, but rather promote a configuration or rearrangement within the dimer that enhances the BRET^2^ signal. Neither NP-001 nor PP-SCR-001 treatment (100 μM) altered the BRET^2^ saturation curve, compared to the vehicle-treated condition, supporting our hypothesis that the actions of PP-001 follow membrane tethering and are sequence-specific.

**Fig. 4.**
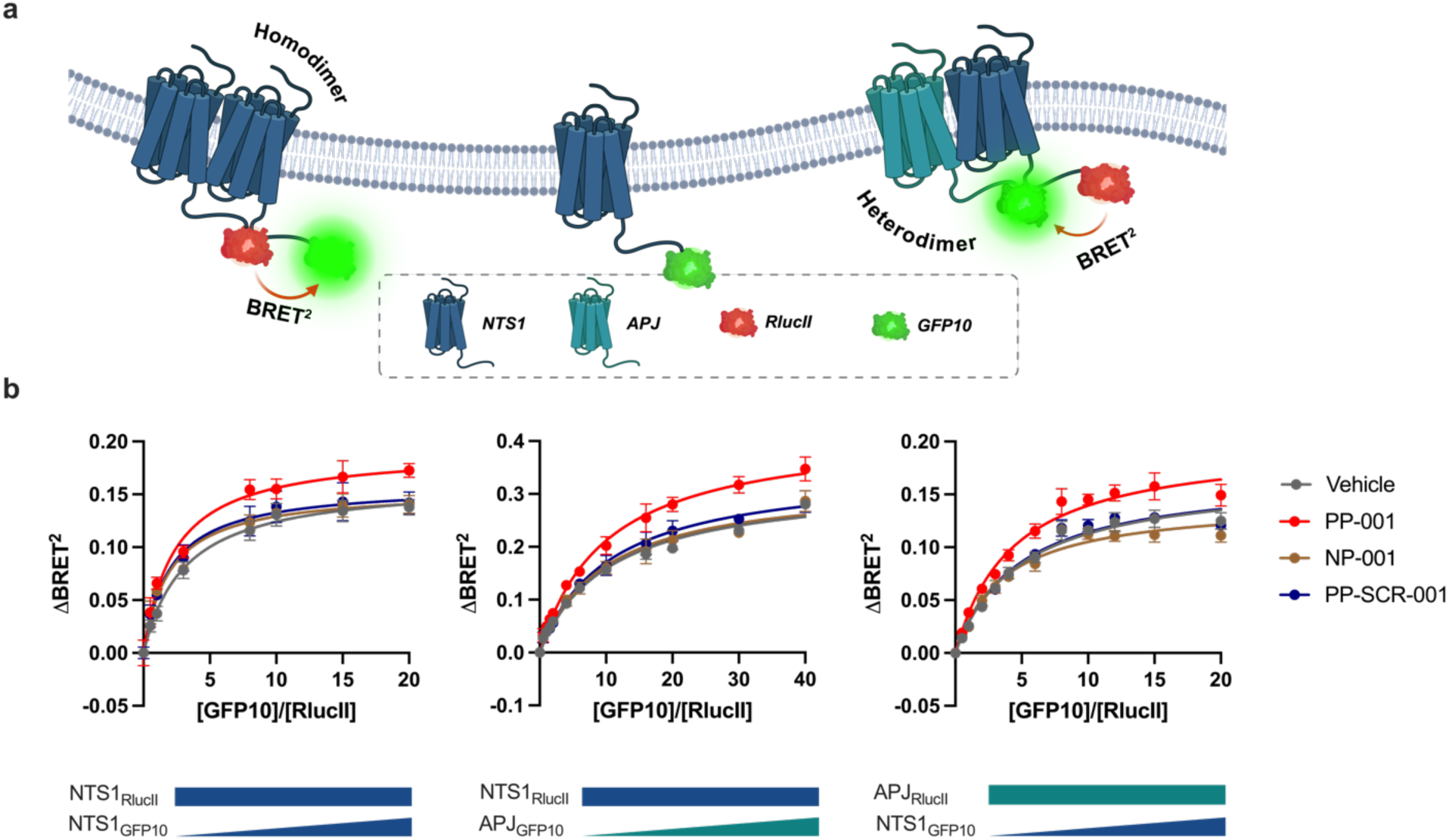
PP-001 promotes receptor-receptor interactions. (a) Principle of BRET^2^ homo- and heteromerization assay. Formation of multimeric donor (RlucII) and acceptor (GFP10)-tagged receptor units leads to an increase in BRET^2^ signal. (b) BRET^2^ titration experiments monitoring receptor-receptor interactions in transiently transfected CHO-K1 (NTS1_RlucII_-NTS1_GFP10_) and HEK293 (NTS1_RlucII_-APJ_GFP10_, APJ_RlucII_-NTS1_GFP10_) cells treated with vehicle (HBSS, 1% DMSO) or pepducin series PP-001, NP-001, and PP-SCR-001 (100 μM). In the NTS1_RlucII_-APJ_GFP10_ experiments (middle panel), cells were treated with 50 μM of pepducin series. In these titration experiments, cells were transfected with fixed amounts of 7TMR_RlucII_ and increasing amounts of 7TMR_GFP10_; specific receptor-receptor interactions is evidenced by a saturation curve rather than a linear increase. Data represent mean ± SEM of three independent experiments, tested in duplicate. Schemas was created with Biorender.com.

In addition to forming NTS1-NTS1 homomers, NTS1 has been reported to form heteromers with several other 7TMRs, including the neurotensin receptor type 2 (NTS2) ^77,80^, the dopamine receptor type 2 (D2R) ^81,82^, the kappa opioid receptor (KOPR) ^83^, and the apelin receptor (APJ) ^84^. Here, we chose to monitor NTS1-APJ interactions by performing similar BRET^2^ titration experiments in HEK293 cells. To increase the robustness of our findings, we performed these experiments using both biosensor pairings: NTS1-RlucII / APJ-GFP10 (**Fig. 4b**, middle panel) and APJ-RlucII / NTS1-GFP10 (**Fig. 4b**, right panel). For one of these pairings (NTS1-RlucII / APJ-GFP10), we reduced the pepducin concentrations to 50 μM, to determine whether this lower concentration was sufficient to enhance heteromerization. As in the NTS1-NTS1 assays, we observed clear saturation curves in cells treated with the vehicle, indicating specific receptor-receptor interactions. Consistent with the homomeric assays, treatment with PP-001 resulted in a significant increase in the BRET_max_ in both assays, representing an increase of 31 ± 3% and 28.0 ± 0.2% for NTS1-APJ and APJ-NTS1, respectively (**Fig. 4b; Supplementary Table S3**). There was no significant change in the BRET_50_ value for PP-001 compared to the vehicle-treated condition, and treatment with NP-001 or PP-SCR-001 did not significantly alter either parameter. Overall, these findings suggest that PP-001 promotes NTS1 receptor homomer- and heteromer-forming processes.

### 3.5. PP-001 induces a sustained drop in blood pressure in rats

NTS1 activation has been associated with various physiological effects that PP-001 may be able to modulate, notably hypotension. Indeed, NT’s depressor effect on rat blood pressure was reported in the very first paper introducing NT ^85^, earning it the “tensin” portion of its name. Importantly, NT’s hypotensive effects are thought to be primarily mediated by NTS1 and not NTS2, as NTS2 is poorly expressed in vascular and cardiac tissues in both rats and humans ^86,87^, and the NTS1-selective antagonist SR48692 has been shown to inhibit the NT-mediated hypotensive response ^88,89^.

To determine the hypotensive potential of these pepducins, we monitored mean arterial blood pressure (MABP) in carotid-artery-cannulated Sprague-Dawley rats, following intravenous injection of NT(8-13) and NTS1-derived pepducins. As shown in **Fig. 5a**, intravenous administration of NT(8-13) (55 nmol/kg, or 0.045 mg/kg) led to a triphasic response, characterized by an immediate initial decline, reaching -29 ± 3 mm Hg 20 s post-injection, a near-complete return to baseline (-7 ± 3 mm Hg at 75 s post-injection), followed by a second, slower decline that stabilized at -28 ± 6 mm Hg by 200 s post-injection. In contrast to NT’s triphasic response, PP-001 at an equimolar dose (55 nmol/kg), after an initial 40-s delay, produced a potent and sustained drop in blood pressure, reaching a minimum of -37 ± 4 mm Hg by 105 s post-injection (**Fig. 5a** and **5b**). A slow upturn ensued, but 300 s post-injection, MABP had stabilized at -25 ± 3 mm Hg. Importantly, neither NP-001 nor PP-SCR-001 induced hypotension in rats: mean maximum ΔMABP values were -7 ± 2 mm Hg and -5 ± 2 mm Hg, respectively, compared to -4.8 ± 0.7 mm Hg for the vehicle-treated control group. Finally, when the NTS1 antagonist SR48692 (1 mg/kg) was pre-administered to rats 10 min prior to PP-001 (55 nmol/kg), the hypotensive effect of PP-001 was significantly reduced (54% reversal), reaching only -17 ± 4 mm Hg, supporting that the in vivo action of PP-001 is mediated by NTS1.

**Fig. 5.**
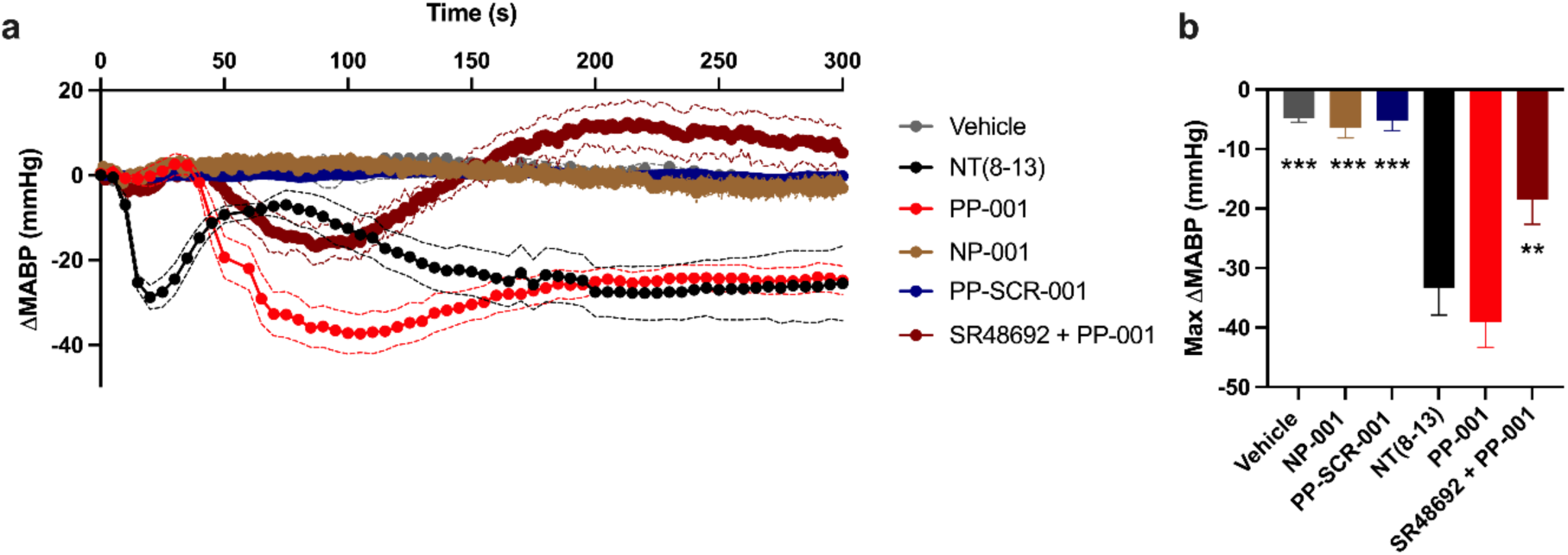
PP-001 induces a sustained drop in arterial blood pressure in rats. (a) Changes in mean arterial blood pressure (MABP) recorded in carotid-artery cannulated male Sprague-Dawley rats over a 5 min-period, following injection into the jugular vein of vehicle (saline, 5% DMSO), NT(8-13), PP-001, NP-001, or PP-SCR-001 at 55 nmol/kg doses. In the SR48692 + PP-001 condition, rats received the NTS1-selective non-peptide antagonist SR48692 (1 mg/kg) 10 minutes prior to PP-001 (55 nmol/kg). (b) Maximal arterial blood pressure decreases (ΔMABP) recorded for each condition. Data represent mean ± SEM, *n* = 4 (NP-001, SR48692 + PP-001), *n* = 5 (vehicle, NT(8-13), PP-SCR-001), *n* = 7 (PP-001) rats. Data were statistically analyzed by a One-way ANOVA followed by Dunnett’s multiple comparisons test, comparing each condition to PP-001-treated condition. ***, *p* < 0.001; **, *p* < 0.01.

### 3.6. PP-001’s N-terminal RKK motif is critical to its cellular actions

To gain mechanistic insight into how PP-001 exerts its effects, and more specifically the relative importance of each amino acid side chain in the PP-001 peptide sequence, we synthesized a 13-pepducin series corresponding to an alanine scan of PP-001. To do so, the residues in positions 2 to 14 of PP-001 ( corresponding to residues R90 to H102 of hNTS1) were sequentially replaced by alanine residues (Note that the residue in position 1 of PP-001, corresponding to A89 of hNTS1, is already an alanine). The sequences of these pepducins are provided in **Table 1**. To characterize the impact of these substitutions on PP-001’s biological activity, we used a label-free technique that measures the global response of a cell monolayer to ligand treatment, namely Electric Cell-substrate Impedance Sensing (ECIS). In this functional assay, cells are seeded onto cell culture plates incorporating gold-plated electrodes at the bottom of each well, through which an alternating electric current is passed. The resistance to the current’s flow both across (transcellular) and between (paracellular) the cells is monitored. The ligand-induced dynamic mass redistribution (DMR) in the cell monolayer, which may include changes in morphology, cytoskeletal organization, and cell-substrate or cell-cell adherence, is accompanied by changes in electrical resistivity ^59,90^.

Here, the DMR kinetics of CHO-K1 cells stably expressing NTS1 were monitored for 1 hour following treatment with 10 μM pepducin. We selected the 10-μM concentration for pepducin testing, having previously found this concentration to be sufficient to promote a cellular response in a similar whole-cell integrated response assay ^42^ and wishing to maintain a low concentration to visualize early differences in the response profile. While the application of PP-SCR-001 did not induce any changes to electrical resistance, cell stimulation with PP-001 triggered an immediate increase that reached a near-plateau by 180 s (3 min), with a maximum change factor of 1.06 ± 0.01 by 1110 s (18.5 min) post-stimulation (see **Fig. 6**). This increase was followed by a gradual decrease that intercepted the x-axis at 2700 s (45 min) post-stimulation. This response profile was similar, but not identical, to the resistivity traces induced by NT and NT(8-13) previously reported ^59^, in which NTS1 activation was also associated with an increase in resistance.

**Fig. 6.**
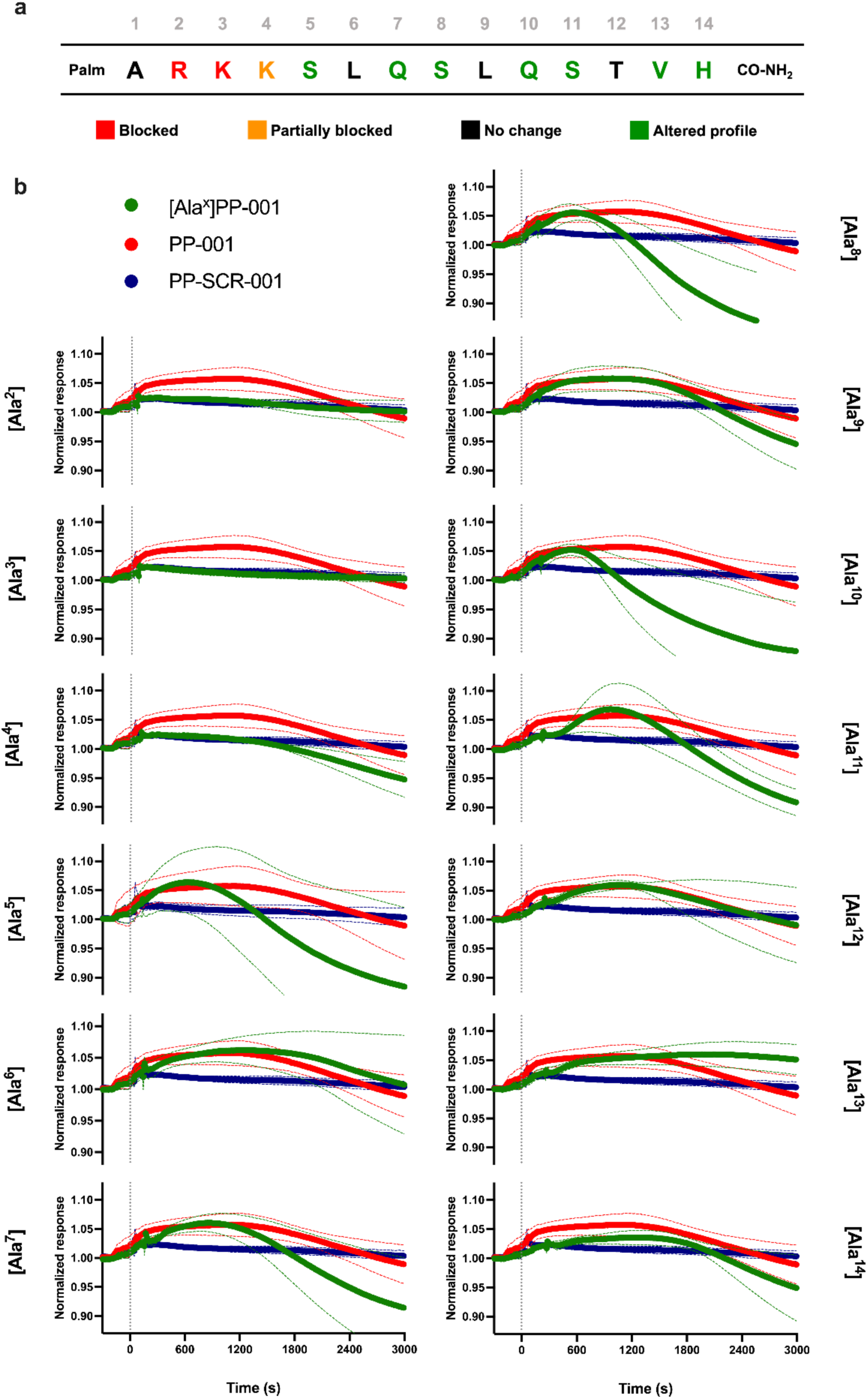
An alanine scan of PP-001 reveals critical residues for biological activity. (a) Color-coded representation of the effects of alanine substitutions on PP-001-induced biological activity. Alanine substitutions of the amino acids shown in red and orange resulted in a total or almost complete loss, respectively, of the pepducin-induced cellular response in an assay monitoring the dynamic mass redistribution (DMR) kinetics of CHO-hNTS1 cells. For those shown in black, a response profile similar to that of PP-001 was observed; for those indicated in green, the response profile was altered. (b) DMR kinetics of CHO-K1 cells stably expressing NTS1, treated with 10 μM concentrations of PP-001, PP-SCR-001, or each of the PP-001 alanine scan pepducins. DMR was monitored by Electric Cell-substrate Impedance Sensing (ECIS) for 50 min post-stimulation. Data represent mean ± SEM, *n* = 3, tested in duplicate. Each condition was normalized to reflect a “normalized response” (or “change factor”) relative to baseline resistance values (ohms)

Subsequently, each pepducin in the alanine scan series was tested, and their profiles were compared to those of PP-001. A color-coded representation of the impact of each substitution on the cellular response induced by PP-001 is provided in **Fig. 6a**, while the complete resistance traces are provided in **Fig. 6b**. As clearly shown, pepducins with substitutions at positions 2 and 3, [Ala^2^]PP-001 and [Ala^3^]PP-001, were completely inactive in this ECIS assay, with their traces being superimposable on those of the negative control PP-SCR-001. [Ala^4^]PP-001 also produced a flat-line response up to 1770 s (29.5 min) post-stimulation, at which point a slow decrease in resistance occurred, culminating in a minimum factor of change value of 0.94 ± 0.04 at 3000 s (50 min) after stimulation. Overall, this suggests that the N-terminal RKK motif is critical for the biological activity mediated by P-001, as changes in these residues completely or quasi-completely abolished the PP-001-induced cellular response. The importance of the RKK motif is further highlighted by the fact that a C-terminally truncated pepducin containing only the peptide sequence ARKKSL (PP-005) displays a signaling signature identical to that of PP-001 (see **Supplementary Fig. S1**). However, PP-005 was less effective than PP-001 in reversing nociceptive behaviors in a tonic pain model (formalin) in rats ^42^, which is why the full-length pepducin was retained for further characterization.

Of the remaining pepducins in the alanine scan series, three produced resistivity traces that were very similar to PP-001: [Ala^6^], [Ala^9^], and [Ala^12^]PP-001. This suggests that L6, L9, and T12 are residues of minor structural importance to PP-001-mediated activity. In contrast, stimulation with the seven other pepducins led to substantially altered response profiles. With the exception of [Ala^14^]PP-001, each produced an increase in amplitude resistance similar to that of PP-001, but with altered kinetics and, in many cases, a more marked decrease below the baseline. [Ala^5^]PP-001, for example, reached its maximum (1.06 ± 0.03) at 600 s (10 min) post-stimulation, and then intercepted the baseline 1475 s (24.6 min) after stimulation, which is much earlier than for PP-001. The subsequent decrease in resistance reached levels of 0.88 ± 0.08 by 3000 s (50 min). [Ala^8^] and [Ala^10^]PP-001 displayed similar profiles to [Ala^5^]PP-001, reaching their maximum values at 543 s (9.1 min) and 536 s (8.9 min) and intercepting the baseline at 1254 s (20.9 min) and 1087 s (18.1 min), respectively. In addition, the [Ala^7^] and [Ala^11^]PP-001 profiles are distinguished by an initial delay in response, as the increase in electrical resistance induced by these pepducins began around 300 s (5 min) and 420 s (7 min) post-treatment, respectively. Each trace reached its maximum (1.06 ± 0.02 and 1.07 ± 0.04) by 880 s (14.7 min) and 979 s (16.3 min) after treatment, respectively. As with [Ala^5^], [Ala^8^], and [Ala^10^]PP-001, the baseline-to-baseline timeframe was much shorter for [Ala^7^] and [Ala^11^]PP-001 than for PP-001, extanding over a period of 25.4-min and 24.1-min period, respectively, compared to 45-min for PP-001. The subsequent decrease in resistance persisted below the baseline (change factors of 0.91 ± 0.9 and 0.91 ± 0.02, respectively, 3000 s post-treatment). As indicated above, the cellular response induced by 10 μM [Ala^14^]PP-001 was the only non-flatline trace that produced a lower amplitude response than PP-001, with a maximum change factor of 1.04 ± 0.01 by 1204 s (19.4 min) post-treatment and a return to baseline by 2227 s (37.1 min). Despite the lower amplitude, the kinetics observed resembled PP-001’s trace. Finally, treatment with [Ala^13^]PP-001 resulted in an increase in resistance that was maintained throughout the readout period. This increase first stabilized around 640 s (10.7 min) and then slowly increased to a maximum of 1.06 ± 0.02 by 1962 s (32.7 min). At the end of the readout (3000 s, or 50 min), the change factor was still above baseline (1.05 ± 0.03). Collectively, these substitutions highlight the importance of the N-terminal RKK motif for the activity of PP-001, although several sites may be of interest for further understanding the mechanisms of action of PP-001.

### 3.7. Alanine substitutions at positions 2 and 13 of PP-001 alter the pepducin’s behavior in vitro and in vivo

Among the above series of pepducins, two were selected for further characterization: [Ala^2^]PP-001 and [Ala^13^]PP-001. In the case of the former, we sought to assess the extent to which the RKK motif is essential to PP-001, in particular whether the inactivity of [Ala^2^]PP-001 observed with the ECIS assay extended to other in vitro and in vivo functional assays. We also selected [Ala^13^]PP-001, as it was the only pepducin with a modified response profile for which the resistance-enhancing effect persisted longer than for PP-001; for all the others, the timeframe was shortened. We therefore sought to determine whether this altered response profile translated into increased potency and/or efficacy. Consequently, both pepducins were subjected to the same battery of tests as those performed for PP-001; the sequences of each pepducin are provided in **Fig. 7a**. Overall, we observed a trend toward complete loss or significant negative shifts in activity for [Ala^2^]PP-001, while [Ala^13^]PP-001 tended to yield similar, sometimes slightly more potent, effects than PP-001 (**Fig. 7b-i**).

**Fig. 7.**
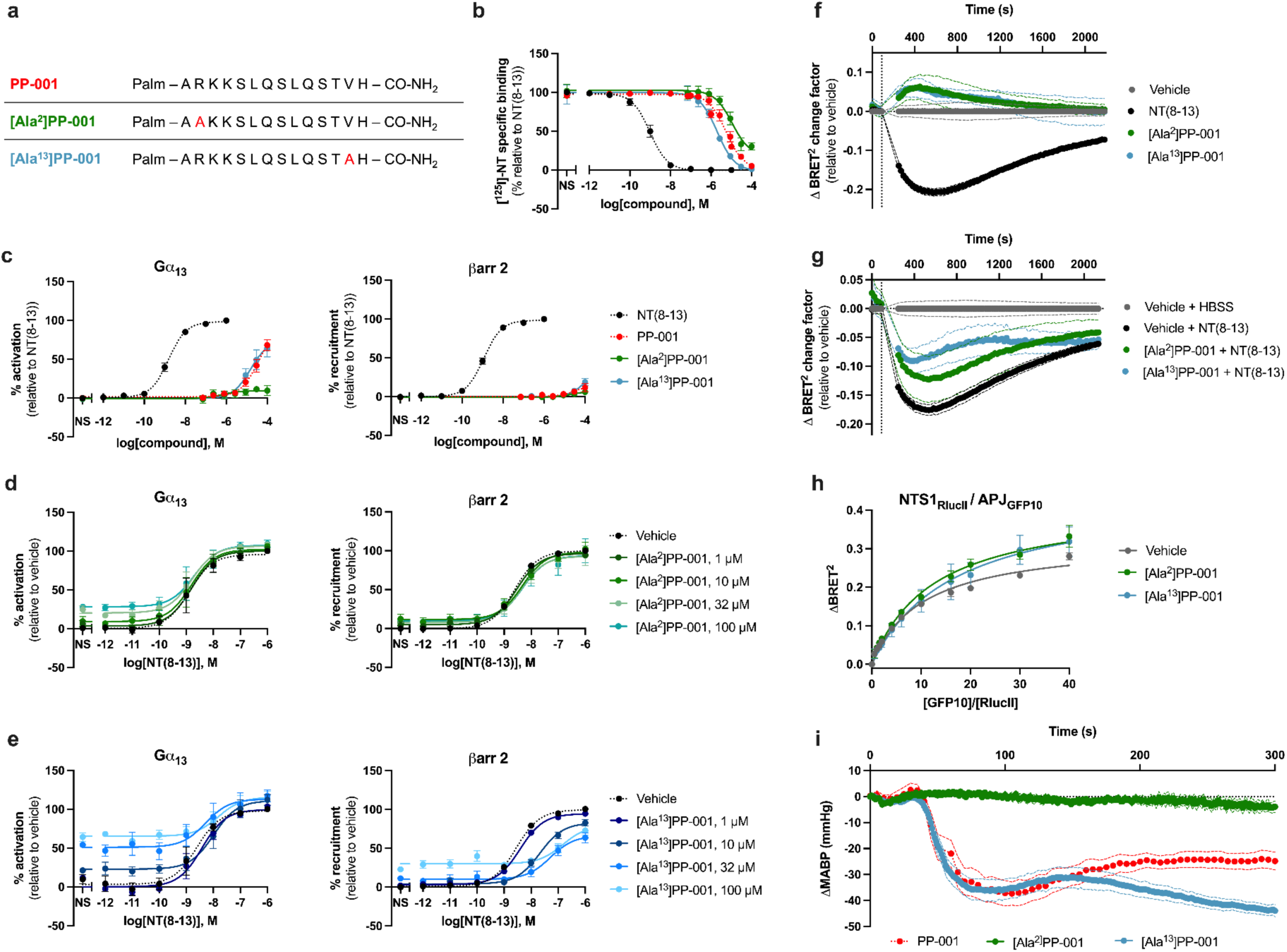
[Ala^2^]PP-001 exhibits reduced biological activity compared to PP-001, whereas [Ala^13^]PP-001 behaves similarly. (a) Table comparing the peptide sequences of PP-001, [Ala^2^]PP-001, and [Ala^13^]PP-001. Alanine substitutions are shown in red. (b) Displacement of ^125^I-radiolabeled NT following incubation of CHO-hNTS1 membranes with increasing concentrations of NT(8-13), PP-001 (for reference), [Ala^2^]PP-001, and [Ala^13^]PP-001. *n* = 3, tested in triplicate. (c) Gα_13_ protein dissociation and βarr 2 recruitment, monitored by BRET^2^, in CHO-K1 cells transiently expressing NTS1, in response to increasing concentrations of NT(8-13), PP-001 (for reference), [Ala^2^]PP-001, and [Ala^13^]PP-001. *n* = 4, tested in triplicate. (d,e) NT(8-13)-induced Gα_13_ protein dissociation and βarr 2 recruitment in CHO-K1 cells transiently expressing NTS1, pre-treated (10 min) with fixed concentrations (0, 1, 10, 32, and 100 μM) of [Ala^2^]PP-001(d) and [Ala^13^]PP-001 (e), as monitored by BRET^2^. *n* = 4, tested in triplicate. (f) NTS1 internalization over a 30-min period, in response to treatment with vehicle (HBSS, 1% DMSO), NT(8-13) (1μM), [Ala^2^]PP-001 (50 μM), or [Ala^13^]PP-001 (50 μM), in transiently transfected CHO-K1 cells. *n* = 4, tested in duplicate. (g) NT(8-13) (1 μM)-induced NTS1 internalization over a 30-min period, in transiently transfected CHO-K1 cells pre-treated (15 min) with vehicle (HBSS, 1% DMSO), [Ala^2^]PP-001 (50 μM) or [Ala^13^]PP-001 (50 μM). *n* = 4, tested in duplicate. (h) NTS1_RlucII_-APJ_GFP10_ interactions in HEK293 cells treated with vehicle, [Ala^2^]PP-001 (50 μM) or [Ala^13^]PP-001 (50 μM), as monitored via BRET^2^ titration experiments. *n* = 3, tested in triplicate. (i) Blood pressure changes over a 5 min period recorded in carotid artery-cannulated male Sprague-Dawley rats, following injection into the jugular vein of PP-001 (for reference), [Ala^2^]PP-001 or [Ala^13^]PP-001 (55 nmol/kg). *n* = 5. Data represent mean ± SEM. BRET^2^ luminescence signaling data. (c, d, e) were recorded 30 min post-pepducin stimulation. NS: non-stimulated (treated with vehicle only).

First of all, in the competitive radioligand binding assay, we observed noticeable negative and positive shifts in potency for the [Ala^2^]PP-001 and [Ala^13^]PP-001 displacement curves, respectively (**Fig. 7b**). Indeed, [Ala^2^]PP-001 displaced radiolabeled NT with an IC_50_ of 11 ± 5 μM, while [Ala^13^]PP-001 exhibited an IC_50_ of 2.1 ± 0.5 μM, contrasting with the IC_50_ of PP-001 of 5.5 ± 0.8 μM. Note that at 100 μM, [Ala^2^]PP-001 did not fully displace radiolabeled NT, but retained 30 ± 8% specific binding.

In BRET^2^ agonist signaling assays (**Fig. 7c**), [Ala^2^]PP-001 led to minimal Gα_13_ protein dissociation (10 ± 7% activation at 100 μM vs 68 ± 7% activation for PP-001), whereas [Ala^13^]PP-001 exhibited allosteric agonist activity similar to that of PP-001 (64 ± 11% activation at 100 μM). Like PP-001, none of the alanine-substituted pepducins triggered significant recruitment of βarr2 to NTS1. In allosteric modulator-type assays (**Fig. 7d-e**, **Supplementary Table S4**), [Ala^2^]PP-001’s inhibitory effect on NT(8-13) signaling was significantly reduced compared to PP-001. Indeed, in the Gα_13_ protein signaling assay, the EC_50_ and E_max_ values of NT(8-13) were not significantly altered by [Ala^2^]PP-001. Likewise, [Ala^2^]PP-001 did not induce any reduction in E_max_ of NT(8-13) in the βarr2 assay (100 ± 20% recruitment at 100 μM [Ala^2^]PP-001 vs 20 ± 20% for PP-001). As with PP-001, pre-treatment of CHO-hNTS1 cells with [Ala^13^]PP-001 resulted in a baseline increase of 64 ± 6% activation in the Gα_13_ assay, consistent with its allosteric agonist action and induced a slight negative shift in EC_50_. In the βarr2 assay, there was a statistically significant shift in both the potency and efficacy of NT(8-13), with a maximum 54 ± 16-fold increase in EC_50_ values and a maximum 36 ± 7% inhibition of the E_max_.

In assays monitoring NTS1 receptor internalization, neither [Ala^2^]PP-001 nor [Ala^13^]PP-001 triggered a decrease in BRET^2^ at 50 μM, as expected given their lack of agonist activity in the βarr2 recruitment assay (**Fig. 7f**). However, both pepducins, but particularly [Ala^13^]PP-001, were able to inhibit NT(8-13)-triggered NTS1 endocytosis (**Fig. 7g**). Indeed, the decrease in the BRET^2^ signal relative to vehicle-treated cells observed after 1 μM NT(8-13) was significantly inhibited by 33 ± 19% and 50 ± 8% by 50 μM [Ala^2^]PP-001 and [Ala^13^]PP-001, respectively. In comparison, pre-treatment with 50 μM PP-001 resulted in a 43 ± 9% inhibition of NTS1 endocytosis. The inhibition mediated by [Ala^2^]PP-001 was unexpected, given the absence of NT(8-13)-mediated βarr2 recruitment inhibition observed in the signaling assays. Nevertheless, the inhibitory effect of [Ala^2^]PP-001 was more moderate than that of PP-001.

In **Fig. 7h**, the effect of 50 μM [Ala^2^]PP-001 and [Ala^13^]PP-001 on NTS1-APJ heteromer interactions was monitored using the hNTS1-RlucII and hAPJ-GFP10 biosensor pairing. In both cases, an upward shift of the BRET^2^ saturation curve was observed. Compared to vehicle-treated cells, treatment with [Ala^2^]PP-001 and [Ala^13^]PP-001 led to a significant increase in BRET_max_, (corresponding to an increase of 17 ± 7% and 14 ± 7%, respectively), which was significantly lower than the 31 ± 3% increase observed with PP-001. The BRET_50_ for each curve was 12 ± 1 and 17 ± 3, respectively, representing a statistically significant increase compared to that of vehicle-treated cells (10.3 ± 0.3) in the case of [Ala^13^]PP-001.

Finally, both pepducins were administered intravenously to Sprague-Dawley rats to determine their hypotensive potential, at equimolar doses to PP-001 (55 nmol/kg) (**Fig. 7i**). Despite the biological activity demonstrated by [Ala^2^]PP-001 in several of the above assays, no change to MABP was observed. In contrast, [Ala^13^]PP-001 induced a potent and sustained drop in blood pressure, initially following the same profile as PP-001, but diverging at the 140-s time-point. The same 40-s delay in response was observed for [Ala^13^]PP-001 as for PP-001, followed by a rapid blood pressure decline, descending to -36 ± 4 mm Hg by 93 s post-administration. There was a brief stabilizing period, with a slow upturn toward baseline that reached -31 ± 4 mm Hg at 140 s. Unlike PP-001, this phase was succeeded by a second slow decline. By the end of the experiment (300 s post-administration), we recorded a ΔMABP of -44 ± 2 mm Hg, which is 19 ± 5 mm Hg lower than that of PP-001 at the same time-point. Altogether, these data highlight the importance of the lysine in position 2 for PP-001 activity and suggest that the valine at position 13 may be an interesting starting point for future structure-activity studies seeking to increase the pepducin’s potency.

### 3.8. PP-001 binds directly to the rat NTS1 receptor

As shown in **Fig. 2**, PP-001 inhibited the specific binding of NT(8-13) and allosterically modulated NTS1 signaling, thereby confirming the pharmacological action of PP-001 on NTS1. To further investigate the mechanism of action of PP-001 on NTS1, we monitored the direct interaction between PP-001 and the receptor in vitro using a N-[4-(7-diethylamino-4-methyl-3-coumarinyl)phenyl]maleimide (CPM)-based stability assay on purified recombinant NTS1 ^64^. This assay has been used to solve the mechanism of ligand binding to membrane proteins ^64,91^. Briefly, Thi CPM stability assay monitors the impact of ligand binding on heat-induced denaturation of a receptor. GPCR purification is a challenge due to their inherent lack of in vitro stability. As such, protein engineering is commonly used to obtain stable 7TMR eluate. We therefore took advantage of two previously described engineered NTS1 constructs (NTS1-3 and NTS1-22) used in structural biology studies (**Supplementary Fig. S2 and Fig. S3**) ^61^ to measure the thermostability of NTS1 in the presence of vehicle, NT(8-13) and PP-001 (**Fig. 8**) (The full list of mutations in the NTS1-22 construct can be found in **Supplementary Table S5**). For the two purified constructs NTS1-3 and NTS1-22, we found that NT(8-13) increased the resistance of NTS1 to heat-induced denaturation, as indicated by the rightward shift of the thermodenaturation curve relative to the vehicle-treated sample (**Fig. 8a-b**). The extent of the shift could be quantified by the melting temperature (Tm), defined as the temperature at the maximum of the curve’s first derivative (**Fig. 8c-d**). We found that NT(8-13) stabilized NTS1 by approximately 6°C for both constructs compared to the vehicle-treated receptor. In contrast, treatment with PP-001 decreased the heat resistance of both NTS1 constructs by approximately 3-4°C, as shown by a left-shift in the thermodenaturation curves. Accordingly, the change in Tm for both NT(8-13) and PP-001 indicates a direct binding of the ligands to purified NTS1. Moreover, the opposite effect of NT(8-13) and PP-001 binding on NTS1 stability suggest that they have a distinct mechanism of ligand action on the receptor, consistent with the allosteric efficacy observed for PP-001 on NTS1-induced NT(8-13) signaling (**Fig. 2b-c**).

**Fig. 8.**
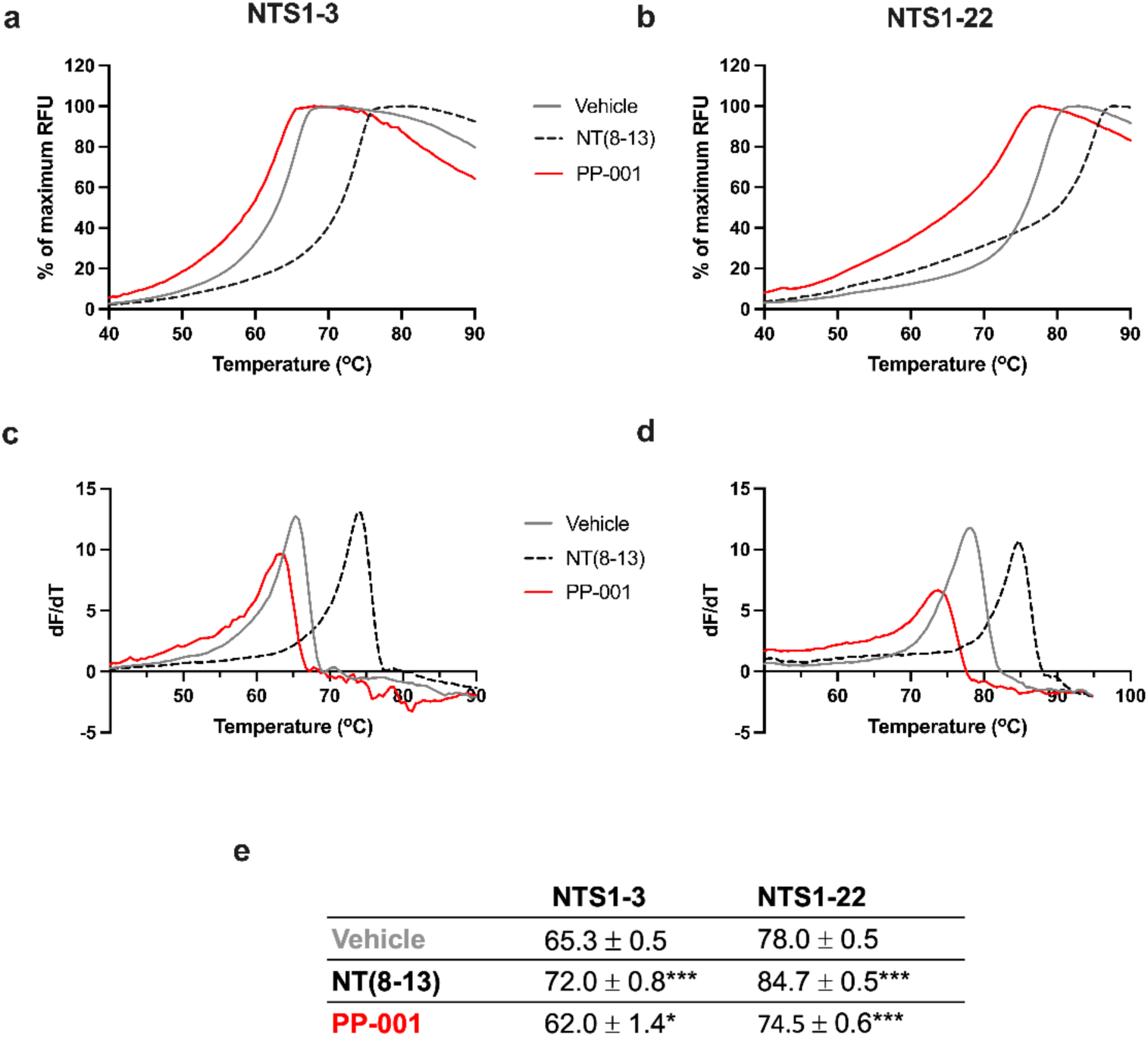
PP-001 binds to NTS1. Representative CPM thermodenaturation curves of purified (a) NTS1-3 or (b) NTS1-22 in the presence of ligands or vehicle. (c, d) First derivative of the CPM thermodenaturation curves presented in (a) and (b), respectively. (e) Table of the Tm values determined from vehicle, NT(8-13) and PP-001 conditions. The data are expressed as a mean ± SEM of 3 independent experiments in duplicate. Statistical significance is determined by a one-way ANOVA followed by a Dunnett’s post-hoc analysis. *** p-value < 0.001; and * p-value < 0.05.

### 3.9. Substitutions of NTS1 ICL residues impede PP-001 activity

Having gathered evidence supporting a direct interaction between pepducin and the receptor, we sought to elucidate the PP-001/NTS1 binding site, with a specific focus on the ICL regions. To do so, we relied on receptor mutagenesis techniques. We introduced a series of sequential alanine mutations into a human HA-tagged NTS1 construct. Our strategy for this first attempt was to “scan” the ICL domains five alanine residues at a time, with the aim of identifying mutants against which PP-001’s activity would be altered. Since the human NTS1 receptor has four ICL sequences spanning 14 (ICL1: A89-H102), 20 (ICL2: E165-R184), 44 (ICL3: A260-R303), and 54 (C-term: N365-Y418) amino acids, as delineated by Uniprot (ID: P30989), 27 mutant constructs were generated (specific sequences can be found in **Supplementary Table S6**). Note that in positions where an alanine was already expressed endogenously, it was simply conserved in the sequence.

We first verified the cell surface expression of the mutants by an enzyme-linked immunosorbent assay (ELISA) performed on transiently transfected HEK293 cells (**Fig. 9a**). Our results revealed that 17 mutants out of 27 were fully expressed, with expression levels greater than or equal to 100% compared to the wild type (WT) receptor. Seven mutants (Mutants 2, 5, 8, 14, 16, 17, and 20) had middling receptor expression levels, ranging from 46 ± 20% (Mutant 8) to 77 ± 20% (Mutant 20). Four were found to be poorly expressed (Mutants 1, 4, 18, and 19), with levels below 25% cell surface expression. Mutant 18 was the least expressed, at 11 ± 5% expression. Thus, in the following assays, the experimental conditions were optimized for each expression group to ensure a valid readout.

**Fig. 9.**
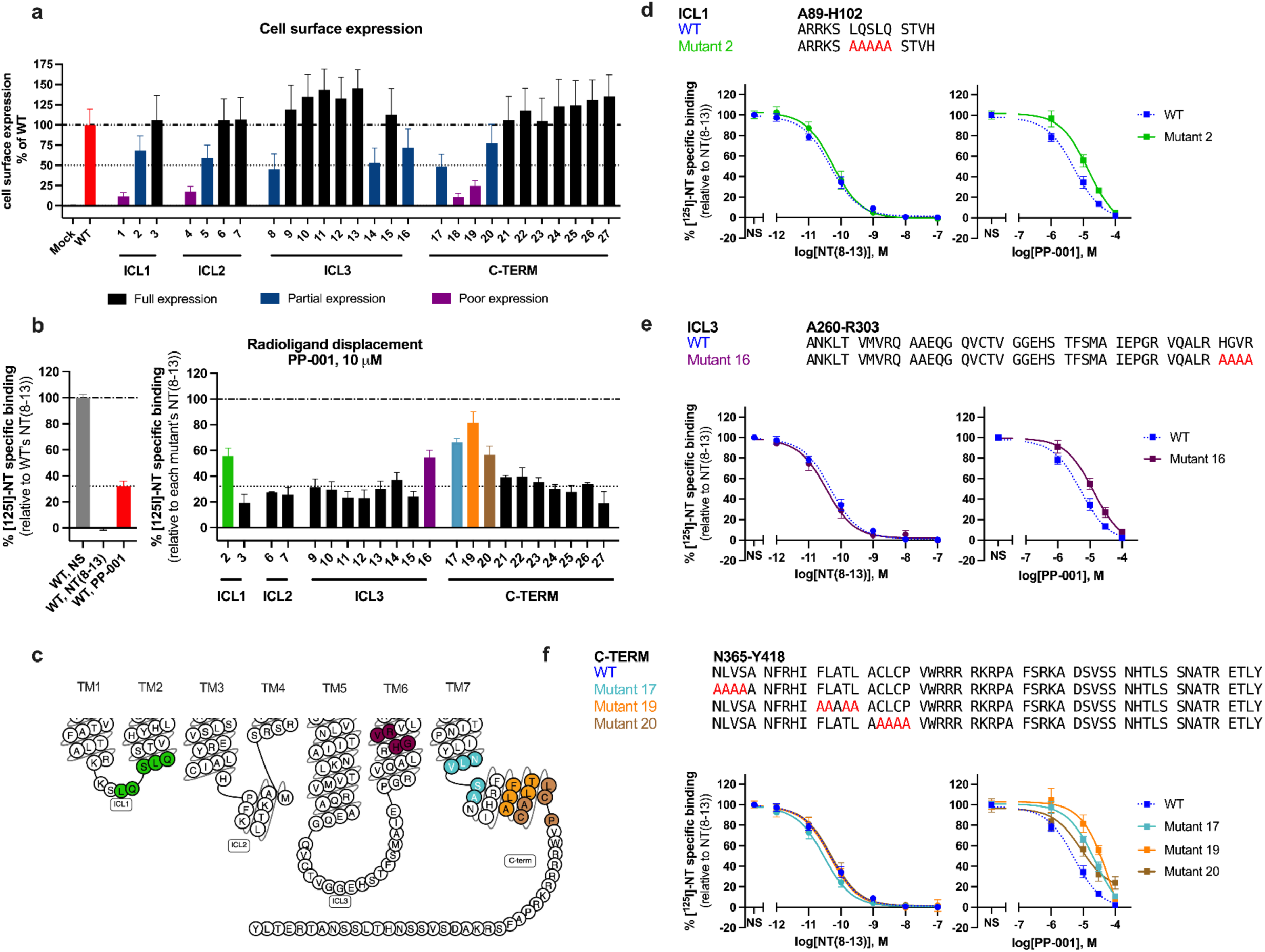
Alanine substitutions in NTS1 ICL domains impede PP-001-driven inhibition of NT binding. (a) Cell surface expression of the HA-hNTS1 WT receptor and the derived ICL mutant constructs (Mutants 1-27) in transiently transfected HEK293 cells, as determined by ELISA. Data was normalized to WT (set as 100% expression) and mock (set as 0% expression) conditions. *n*=3, tested in duplicate. (b) Displacement of ^125^I-radiolabeled NT by 10 μM PP-001 in HEK293 cell membranes transiently transfected with the HA-hNTS1 WT receptor (left) or the ICL mutant constructs (right). No reliable results could be generated with mutants 1, 4, 5, 8, and 18. Mutants represented in color (2, 16, 17, 19, 20) were found to retain greater binding of [^125^I]-NT. *n*=2, tested in duplicate. (c) Snake plot of the human NTS1 receptor, cropped to ICL domains. Substituted residues in Mutants 2, 16, 17, 19 and 20 are identified in color. (d-f) Displacement of ^125^I-radiolabeled NT following incubation of transiently transfected HEK293 membranes with increasing concentrations of “cold” NT(8-13) (left) or PP-001 (right). Data for the WT condition are compared to mutants in the ICL1 (Mutant 2, d), ICL3 (Mutant 16, e) and C-terminal (Mutants 17, 19, and 20, f) regions. Sequences of the ICL domains with relevant mutations are presented above each graph; alanine residue substitutions are shown in red. *n*=2, tested in duplicate. Data represent mean ± SEM. Radioactivity data were normalized according to NT(8-13) (set as 100% specific binding) and untreated cells (set as 0% specific binding) for each individual mutant.

Here, we chose to test the WT receptor and the 27 mutants in the radioligand binding assay. The aim was to compare the inhibitory activity of PP-001 on the orthosteric NT binding at each mutant (relative to NT(8-13)) with that of the WT receptor. **Fig. 9b** shows the results obtained at 10 μM PP-001. For 22 of the 27 mutants, PP-001 was found to inhibit [^125^I]-NT binding to a similar extent as the WT condition. Interestingly, in the case of Mutants 2, 16, 17, 19, and 20, a greater amount of bound radioactivity was retained; in other words, the inhibitory effect of PP-001 on NT binding was partially reversed. In contrast to the 32 ± 4% specific binding retained at the WT receptor, the retained specific binding levels were 56 ± 6% (Mutant 2), 55 ± 5% (Mutant 16), 66 ± 3% (Mutant 17), 82 ± 9% (Mutant 19), and 57 ± 7% (Mutant 20). The locations of these mutations on NTS1 are presented in the snake plot in **Fig. 9c**; they correspond to alanine substitutions in the ICL1 (Mutant 2), ICL3 (Mutant 16), and C-terminal tail (Mutants 17, 19, and 20) sequences of NTS1. It should be noted that Mutants 19 and 20 are found within the H8 domain, while Mutant 17 is adjacent to H8. Unfortunately, no reliable data could be obtained for the 5 remaining mutants (1, 4, 5, 8, and 18).

In **Fig. 9d-f**, the full concentration-response curve data for both NT(8-13) and PP-001 are shown for the five highlighted mutants, along with the specific mutation sequences. The curves for the other 17 mutants are presented in **Supplementary Fig. S4**. The IC_50_ and maximum inhibition values extracted from these curves can be found in **Supplementary Table S7**. As can be seen, while the NT(8-13) curves for each mutant are superimposable on those for WT, the PP-001 curves are slightly but consistently shifted to the right. For Mutants 2 and 16, this represents a nearly three-fold shift in IC_50_: 14 ± 2 μM and 13 ± 2 μM, respectively, compared to 5.3 ± 0.5 μM for the WT receptor. For Mutant 17, this shift is slightly more pronounced (23 ± 3 μM). However, the greatest shift in affinity was observed for Mutant 19, with a shift of 70 ± 30 μM. Interestingly, in the case of Mutant 20, the rightward shift of the PP-001 curve was more subtle (9 ± 3 μM), but PP-001’s inhibition of NT binding at 100 μM was no longer complete. Instead, 24 ± 6% of the specific [^125^I]-NT binding was still retained. Together, these data suggest that the PP-001/NTS1 binding site may incorporate several ICL regions, but that H8 in the C-terminal domain could be a particularly interesting area.

## 4. Discussion

In the present study, we sought to characterize the modulatory effects of a pepducin derived from the first intracellular domain of NTS1, namely PP-001, on target receptor function and to provide important insights into the mechanisms by which these pepducins exert their modulating action. Our results revealed that this lipidated cell-penetrating peptide, acting as a biased allosteric agonist, selectively activates the downstream G-protein signaling pathways, favors homomer and heteromer formation, and is highly effective in lowering blood pressure. PP-001 was also found to behave as a negative allosteric modulator of NTS1, decreasing the receptor’s ability to bind to its natural orthosteric ligand and inhibiting NT-promoted βarr recruitment and NTS1 receptor internalization. Finally, using alanine scanning experiments and purified NTS1 receptor constructs, we also identified a critical N-terminal motif in the pepducin sequence for the biological function of PP-001 and showed that the eighth α-helix (H8) of NTS1 represents an important site at the interface of pepducin-receptor interaction.

Through a series of in vitro assays, we accumulated evidence suggesting that PP-001 preferentially activates G protein signaling pathways rather than βarr pathways, while simultaneously inhibiting NT binding. The pepducin-induced loss of NT binding was in itself an uncommon result, as several research groups have described pepducins that have no to little impact on the orthosteric binding of endogenous ligands to their target receptors. These include β2AR-derived pepducins ICL1-9 ^7^, ICL3-8, and ICL3-9 ^6^ (determined by competitive radioligand binding experiments) and the CXCR4-derived pepducin ATI-2341 ^4^ (determined by competitive TagLite FRET assays). The PAR4-derived pepducin P4pal-10 inhibited FPR2-driven responses in neutrophils without directly competing for ligand binding (determined by flow cytometry with a fluorescently labeled FPR2 agonist) ^92^. To our knowledge, the only pepducins reported to “compete” for binding with an extracellular agonist are the pepducins F1pal-16 and F2pal-10 targeting FPR2 (determined by flow cytometry) ^19–21^. In these publications and others ^93^, the authors raise the question of whether there could be direct pepducin/agonist competition at the extracellular site. Although we cannot categorically rule out this possibility, we interpret this result as the consequence of intracellular allosteric action, particularly in light of the complete lack of effect on NT binding observed with the non-palmitoylated peptide, NP-001. Rather, we believe that the pepducin, receptor, and extracellular ligand form an allosteric system in which they act respectively as modulator, conduit, and guest [to use Kenakin’s terminology ^94^], where the binding of PP-001 induces a conformational change in the receptor that alters NT(8-13) activity and, reciprocally, the binding of NT(8-13) (and potentially the binding of the NTS1 antagonist, SR48692) induces a conformational change that alters PP-001 activity.

Our results further demonstrate that PP-001 simultaneously displays agonist and NAM effects on NTS1 signaling, which is also unusual in pepducin research, although not unprecedented. For example, the urotensin receptor (UTR)-derived pepducins hUT-Pep2 and [Trp^1^, Leu^2^]hUT-Pep3 effectively activate Gα_i_ and Gα_13_ signaling pathways in HEK293 cells while triggering only low levels of Gα_q_ and Gα_12_ activation or βarr recruitment and, importantly, inhibiting urotensin II-driven Gα_12_ signaling and aortic ring contractions ^24^. The β2AR-derived pepducins ICL3-9, as described above, is another example. These observations highlight the complex signaling patterns that may be alternately preserved or induced in a system by allosteric modulators acting selectively on specific signaling pathways ^95^.

The subsequent lack of pepducin-induced NTS1 endocytosis aligns with the G protein-biased signaling profile of PP-001. Indeed, unlike NT(8-13), which promotes receptor internalization (as indicated by a decrease in the BRET^2^ signal), both PP-001 and the control pepducin PP-SCR-001, in agonist mode, produced only a slight, transient, and highly variable increase in BRET^2^ signal. We interpret this as reflecting the pepducin’s insertion into the cell membrane; conceivably, the resulting rearrangement of membrane phospholipids may increase the proximity between receptor and plasma membrane marker or promote a donor/acceptor orientation that favors BRET^2^ signal. In antagonist mode, the inhibition of NT(8-13)-induced NTS1 endocytosis resembles the findings obtained with the Gα_s_-biased pepducin ICL3-9 ^6^, which mimics the ICL3 sequence of β2AR. This pepducin has been shown to enhance cAMP production and Gα_s_/β2AR engagement in HEK293 cells with minimal βarr recruitment, while also inhibiting isoproterenol-promoted βarr recruitment and β2AR internalization. A key difference here is that pepducin ICL3-9 did not affect the orthosteric binding of radiolabeled cyanopindolol to β2AR. Interestingly, these findings also raise the possibility that by preventing agonist-induced 7TMR internalization, such pepducins may retain receptors at the cell surface, threreby further favoring signaling toward G protein pathways.

Collectively, these data suggest that PP-001 has dual modulatory effects on NTS1: PP-001 induces a conformation of the NTS1 receptor that (1) promotes G protein signaling over βarr recruitment (biased allosteric agonist behavior) and (2) impedes NT binding and its subsequent effects (NAM behavior), possibly due to reduced NTS1 thermodynamic stability. However, one difficulty remains: determining whether the ensuing effects include inhibition of NT-mediated G protein-dependent signaling pathways, as would be expected following a reduction in NT binding. Unfortunately, our G protein signaling assays do not provide a conclusive answer. We did not observe any inhibition of NT-induced G protein signaling by PP-001, but this could be effectively masked by the pepducin’s own agonist behavior and by the ability of PP-001 to prevent NTS1 endocytosis. In the absence of additional evidence, we favor the interpretation that all NT-induced signaling pathways, G protein-dependent and G protein-independent alike, are inhibited by PP-001’s NAM behavior.

In addition, we demonstrated for the first time that these pepducins can promote oligomer formation, with PP-001 inducing the formation of NTS1-NTS1 homomers and NTS1-APJ heteromers. The CXCR4-derived, G protein-biased agonist pepducin ATI-2341 was tested in a similar BRET assay but had no effect on receptor homomerization ^13^. This is not to say that the concept of dimerization has not been of interest in pepducin research; indeed, since the very first publication on pepducins in 2002 by Covic *et al.,* the possibility that pepducins actually reflect a protomer-protomer interaction has been raised ^1^. If pepducins do form “pseudo-dimers” with their target receptor, they may be tapping into signaling features that are unavailable to monomeric receptor units. In this specific case, it is unclear whether PP-001 in fact promotes the formation of new NTS1 dimers (creating a pseudo-trimer?) or rather induces a conformational rearrangement within an existing dimer that enhances the BRET^2^ signal. In either instance, our findings suggest a direct receptor-pepducin interaction and highlight another feature of pepducin behavior that may differ from that of extracellular ligands.

Importantly, we also found that this pepducin targeting NTS1 is active in vivo. Indeed, PP-001 induced a potent and sustained drop in blood pressure when administered to rats, an effect consistent with NTS1 activation (or allosteric agonist behavior) ^96^. As observed previously ^97^, intravenous injection of NT(8-13) induced a significant hypotensive effect, characterized by a triphasic response. A rapid drop in blood pressure of -30 mm Hg occurred quickly (first phase) followed by a rapid return to baseline levels (second phase), then a prolonged decrease that stabilized around -30 mm Hg (third phase). Mechanistically, the initial decline in blood pressure is thought to be triggered by NT-mediated mast cell degranulation and histamine release, which in turn triggers the release of catecholamines to counteract the drop in blood pressure ^98^. In contrast, the hypotension induced by PP-001 was sustained and more pronounced than that of NT(8-13) at an equimolar dose. Given the disparity in potency between NT(8-13) and PP-001, the intensity of this response was somewhat unexpected. If the depressive response is mediated by activation of G protein signaling pathways at NTS1, as suggested by the biased signaling profile of PP-001, it is possible that by also inhibiting endogenous NT βarr signaling and NTS1 internalization, PP-001 creates a strong bias that potentiates the hypotensive response. Alternatively, a factor to consider here may be the kinetics of pepducins. Unlike NT(8-13), for which a bolus injection could trigger immediate activation of NTS1 and thus rapid release of histamine, PP-001 must first insert itself into the cell plasma membrane. It is therefore likely that NTS1 activation is a slower and more protracted process. Consequently, histamine release may be delayed and/or staggered over time to create a longer-lasting response that catecholamines may be unable to reverse. Finally, additional features of PP-001, such as the absence of receptor desensitization or the promotion/targeting of multimeric receptor units, may come into play. Importantly, we also demonstrated that the antagonist SR48692, which binds to NTS1 in distinct but overlapping binding domains with NT, blocked the hypotensive action of PP-001, further supporting that pepducin’s effect was mediated by NTS1. Interestingly, a small-molecule, SBI-553, also identified as a NTS1 allosteric modulator, has been described to act as a βarr-biased allosteric agonist, selectively inhibiting NT-induced G protein signaling and behaving as a PAM for βarrs, thus displaying a signaling profile diametrically opposite to that of PP-001 ^99^. Described by cryo-EM as interacting with the receptor’s intracellular cavity ^100,101^, SBI-553 did not to produce a hypotensive response in mice, consistent with our own observation of potent PP-001-induced hypotension and the contrasting biases of SBI-553 and PP-001 ^102^. Overall, these results reinforce the concept of allostery, whereby intracellular modulators such as pepducins influence the binding of orthosteric ligands, but the action of orthosteric agonists or antagonists can in turn modulate the activity of intracellular allosteric ligands.

In addition, we present here a new pepducin alanine scan series that identified a critical N-terminal RKK motif, whose importance was reinforced by the observation that the R2A substitution (i.e., [Ala^2^]PP-001) had a negative impact on the activity of PP-001 in a panel of in vitro and in vivo functional assays. Interestingly, several pepducin studies have highlighted the importance of positively charged residues such as R and K for pepducin activity. For example, Forsman *et al.* reported complete inactivity of the FPR2-derived pepducin F2pal_10_ when lysine at position 5 (K5) is mutated to A, N, or Q, but a return to activity when replaced by a positively charged arginine residue ^20^. Similarly, the activity of P1pal-19 (a PAR1 mimic) was impaired following substitutions at R16 in particular, but also at R11 and K13. For pepducins, it is often unclear whether the main contribution of key amino acids is to facilitate the pepducin flip-flop process, to interact with residues on the receptor (and/or its effectors) to ensure binding, or to fulfill a combination of both functions. In this case, we consider it unlikely that the positively charged lysine residues promote membrane translocation ^103^, and therefore assume that they play a role in PP-001/NTS1 affinity. However, given the multiple residues whose alanine substitutions resulted in altered impedance response profiles, it is clear that deciphering the relationship between the structure of PP-001 and its complex NTS1 modulation pattern remains a challenging task.

Finally, we used a thermal denaturation assay to demonstrate the direct binding between PP-001 and purified NTS1 protein. Although only a few studies have successfully shown pepducin/receptor binding, these studies used various techniques: Janz *et al.* utilized a photochemical cross-linking approach to reveal the direct binding between CXCR4 and ATI-2341 ^4^, Carr *et al.* used a receptor labeled with an environment-sensitive fluorophore (monobromobimane) to monitor conformational changes induced by pepducins on β2AR ^6^, and Zhang *et al.* used biotinylated peptides to immunoprecipitate the target receptor PAR1 ^104^. Here, we show that PP-001 significantly alters the Tm of the NTS1-3 and NTS1-22 constructs and that, intriguingly, this value is left-shifted. Although the opposite effects observed with NT(8-13) and PP-001 are consistent with some observations made in this study, it is interesting to consider that PP-001 might in fact induce less thermodynamically stable receptor conformations. While we are uncertain about the implications of this finding, it does demonstraste a direct pepducin-receptor interaction. To elucidate the site where PP-001 could bind to NTS1, we then generated a series of NTS1 receptor constructs with sequential alanine mutations in the ICL1-3 and C-terminal domains and tested them against NT(8-13) and PP-001 in a radioligand binding assay. While PP-001’s ability to inhibit NT binding was largely retained across the mutant series, its potency was right-shifted for select mutants with substitutions in the ICL1, ICL3, and C-terminal domains, two of which encompass the H8 domain of NTS1 and another adjacent to H8. This is particularly interesting in light of the findings of Zhang *et al*. ^104^, whose 2015 study represents perhaps the most extensive investigation to date of the mode of action of pepducins. They proposed a mechanism in which the allosteric agonist pepducin P1pal-19, derived from the ICL3 sequence of PAR1, would bind to H8 in a manner parallel to the second protomer in a PAR1-PAR1 dimer. Specifically, they showed by immunoprecipitation that direct binding between a biotinylated ICL3 sequence and PAR1 was lost following deletion of the H8 domain. They also performed mutagenesis with targeted mutations in all ICL sequences of PAR1 and, similarly to this study, found that the most negative changes in pepducin activity occurred when H8 residues were mutated. Furthermore, Sevigny *et al*. used a dimeric model of PAR2 featuring ICL3 and H8 binding interactions to design a series of of antagonist pepducins, with great efficacy ^9^. There are, of course, several key differences between PP-001 and P1pal-19, including the fact that they are derived from different ICL domains (ICL1 and ICL3, respectively) of their target receptor. Also, P1pal-19 has been described simply as an allosteric agonist, whereas PP-001 additionally displays signaling bias and NAM effects on NT-induced signaling. The activity of P1pal-19 on select mutants was determined using an agonist assay (measuring inositol phosphate production), while we monitored PP-001’s ability to inhibit NT binding. Despite these differences, the observation that both pepducins may interact with the H8 domains of their respective receptors to regulate 7TMR function suggests that this may be a common mechanism for this unique class of modulators. It is interesting to note that conformational changes in the H8 domain have been shown to govern the biased activation of the mu-opioid receptor (MOR) toward G proteins ^105^, as demonstrated here by PP-001.

## 5. Limitations of this study

An important qualification should be considered. While the thermostability assay was performed with rat NTS1-3 and NTS1-22 receptor constructs, the ICL mutant series was derived from the human NTS1 receptor. Although rat and human NTS1 receptors share 84% sequence identity and the ICL1 sequence is identical between the two species, there may nevertheless be differences in the binding mode between PP-001/rNTS1 and PP-001/hNTS1. Also, it is important to note that while the rNTS1-3 construct has no mutations in the regions corresponding to the substitutions in Mutants 2, 16, 17, 19, and 20 (all relevant sites are intact to allow PP-001/NTS1-3 binding), this is not the case for the rNTS1-22 construct. In fact, the first amino acid substituted in the sequence of Mutant 16 (H305 in rNTS1) is mutated to an arginine, while the last two residues substituted in Mutant 17 (V372 and S373 in rNTS1) are lost in the deletion V372 to Y424, as are the entire substituted sequences in Mutants 19 and 20. Intriguingly, despite the loss of these last three sites, direct binding between PP-001 and rNTS1-22 is preserved. Thus, although our radioligand binding data suggest that H8 is the primary zone of interest for future mechanistic studies, the PP-001 binding site may incorporate residues from multiple intracellular domains. These could be found in ICL1 and ICL3, as suggested by Mutants 2 and 16, or in the sites corresponding to Mutants 1, 4, 5, 8, or 18, for which we were unable to generate data. It is also possible that our strategy of sequentially substituting ICL residues of five amino acids at a time is not an optimal framework for our alanine scan; we may have overlooked important residues that straddle two different mutant sequences. The idea that H8 may not be the only site of interest for a potential PP-001/NTS1 binding site is further highlighted by the fact that, even for Mutant 19, where the potency of PP-001 was right-shifted to the greatest extent among all mutants, there was still complete inhibition of NT binding at 100 μM. This suggests that PP-001 binding was still partially conserved.

In summary, our results provide evidence of the complex interaction between pepducin PP-001 and its target receptor, NTS1, acting simultaneously as a biased intracellular allosteric agonist and as a negative allosteric modulator of NT function. We contend that NTS1-derived pepducins, due to their access to a unique and complex signaling landscape, represent a valuable pharmacological tool in drug discovery that may inform the development of novel therapeutics.

## Acknowledgements

The authors would like to thank Drs. M. Bouvier, T. Hébert, S.A. Laporte, G. Pineyro, R. Leduc, J.-C. Tardif, and E. Thorin for providing us with BRET-based biosensors. R.L.B. is the recipient of Ph.D. scholarships from the Fonds de recherche du Québec – Santé (FRQ-S) and the Canadian Institutes of Health Research (CIHR). F.L. is a recipient of a Master’s-level scholarship from the CIHR, É.B. is a recipient of Master’s-level scholarships from the FRQ-S and the CIHR. É.B.O. is the recipient of an Excellence Research Chair in Innovative Theranostic Approaches in Ovarian Cancers (THERANOVCA) funded by the Région Normandie and cofinanced by the European Union. PS is the recipient of a Tier 1 Canada Research Chair in Neurophysiopharmacology of Chronic Pain and a member of the FRQ-S-funded Québec Pain Research Network (QPRN).

## Author contributions

Conceptualization: C.E.M., E.B.O., P.S.

Formal analysis: R.L.B., V.B.

Funding acquisition: C.E.M., E.B.O., M.G., P.S.

Investigation: R.L.B., E.B., F.L., N.M., M.C., J.C., V.B.

Methodology: R.L.B, C.E.M., E.B.O.

Project administration: E.B.O., J.M.L, M.A., M.G., P.S.

Resources: E.M.

Supervision: P.S.

Visualization: R.L.B., V.B.

Writing – original draft, R.L.B.

Writing – review & editing, J.M.L., M.G., E.B.O., and P.S.

## Data and materials availability

The BRET^2^ constructs Gα_q_-RlucII, Gα_oA_-RlucII, Gα_13_-RlucII, GFP10-Gγ_1_, RlucII-β-arrestin-1, RlucII-β-arrestin-2, and rGFP-CAAX were available to us through a materials transfer agreement (MTA) with Dr. Michel Bouvier’s laboratory (Université de Montréal). All data are available in the main text or in the supplementary materials.

