## Supplementary Information for "A neurotensin receptor type 1-derived pepducin acts as a biased allosteric modulator to regulate target receptor function"

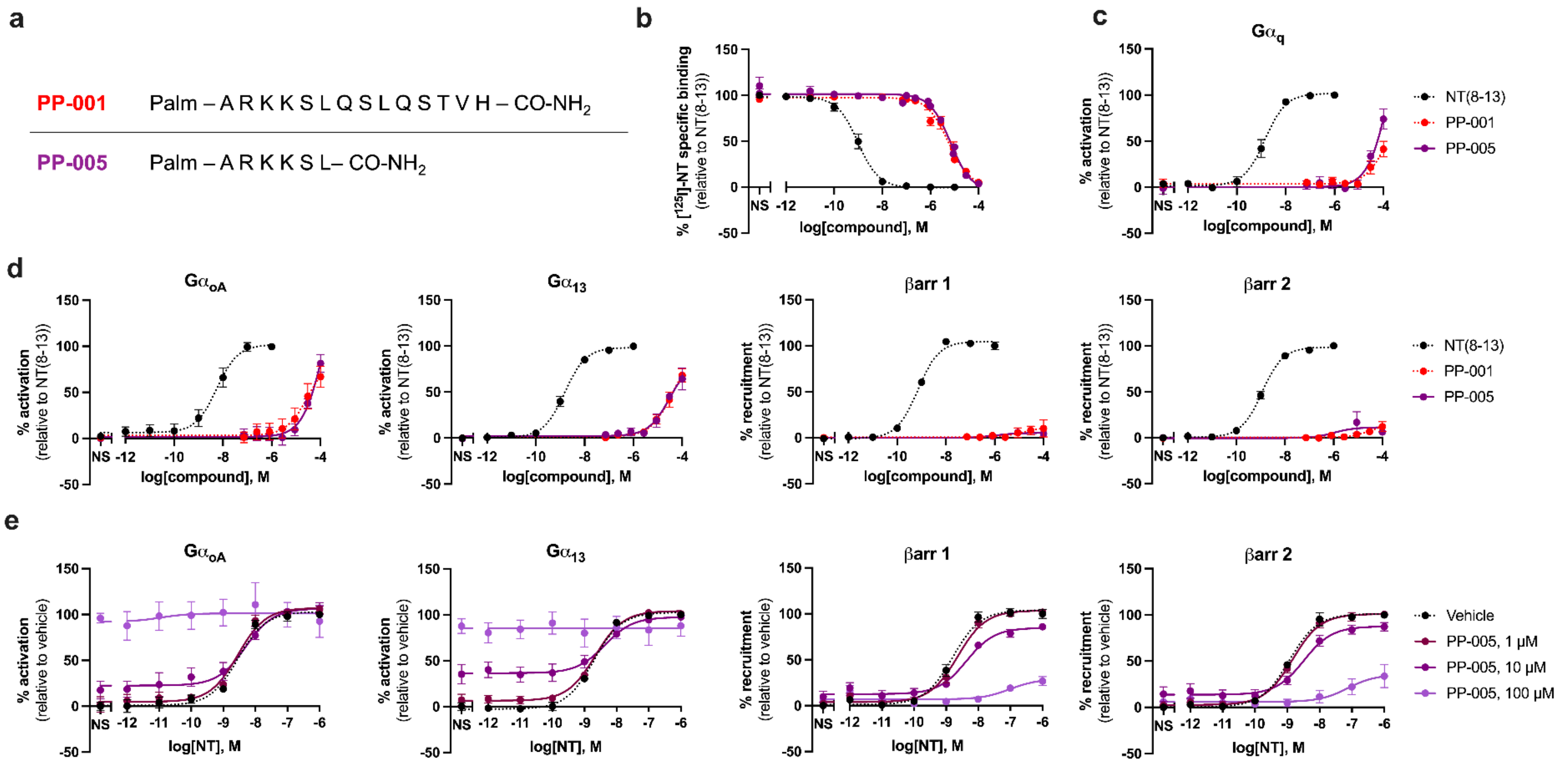


**Supplementary Fig. S1. A C-terminally truncated analog of PP-001 displays a similar signaling signature.** (a) Table comparing the peptide sequences of PP-001 and the C-terminally truncated pepducin PP-005. (b) Displacement of $^{125}$I-radiolabeled NT following incubation of CHO-hNTS1 membranes with increasing concentrations of NT(8-13), PP-001 (for reference), and PP-005. (c, d) G protein dissociation and βarr recruitment in transiently transfected CHO-K1 cells, monitored by BRET$^2$, in response to increasing concentrations of NT(8-13), PP-001, and PP-005. (e) NT-induced G protein dissociation and βarr 2 recruitment in transiently transfected CHO-K1 cells pre-treated (10 min) with increasing concentrations (0, 1, 10, and 100 μM) of PP-005. Data represent mean ± SEM of three independent experiments, tested in triplicate. $BRET^2$ luminescence was recorded 30 min post pepducin stimulation, 20 min post NT(8-13) stimulation. Data were normalized according to NT(8-13): values from untreated conditions were set as 0% specific binding (activation, recruitment) and values from NT(8-13) 1 μM-treated conditions were set as 100 % specific binding (activation, recruitment). NS: non-stimulated (treated with vehicle only.)

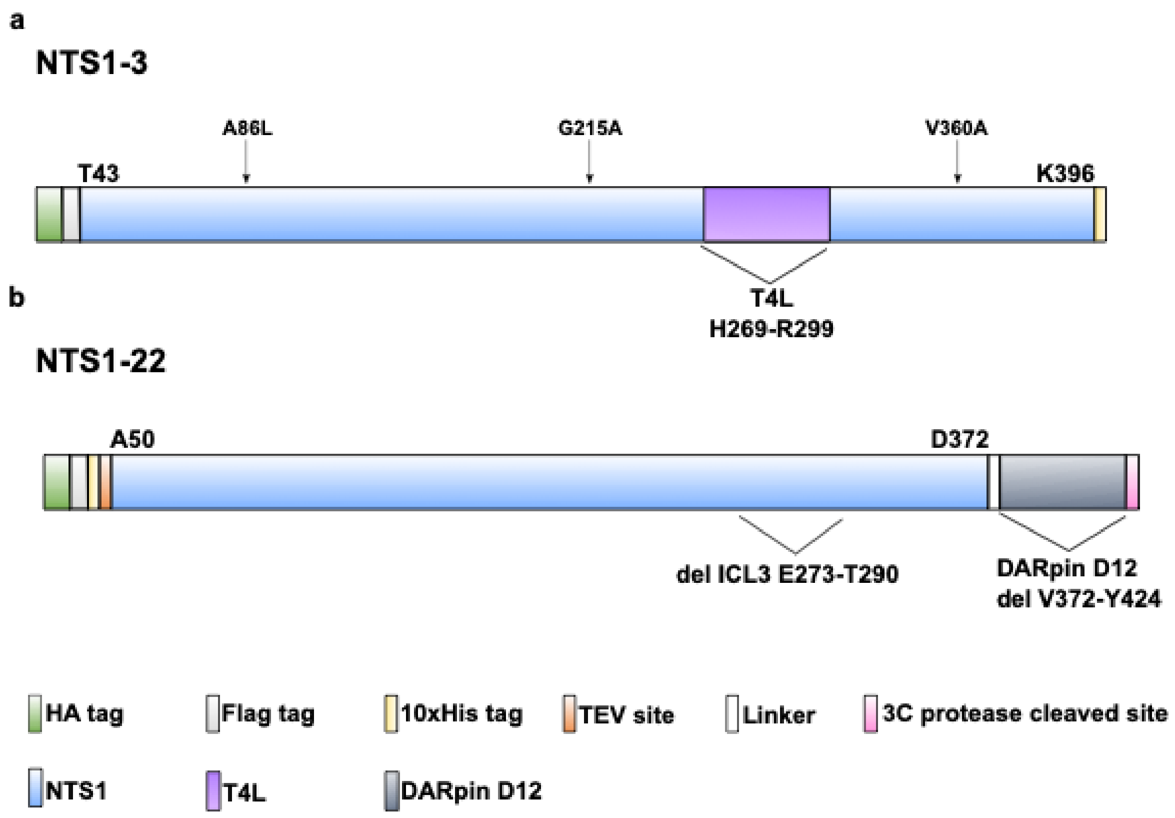


**Supplementary Fig. S2. Schematic representation of NTS1 fusion proteins.** (a) Modifications of NTS1-3 and (b) NTS1-22 proteins. Added tags and soluble domains are represented by colored defined rectangles. N- and C-terminal deletions are identified by the first and last amino acids. Substitutions are delimited by brackets with the amino acid position marking the replaced part. Point mutations are indicated by arrows on NTS1-3 (a) and NTS1-22 (b). NTS1-22 has been named NTS1-H4bm elsewhere[81]. B-W: Ballesteros-Weinstein numbering system.

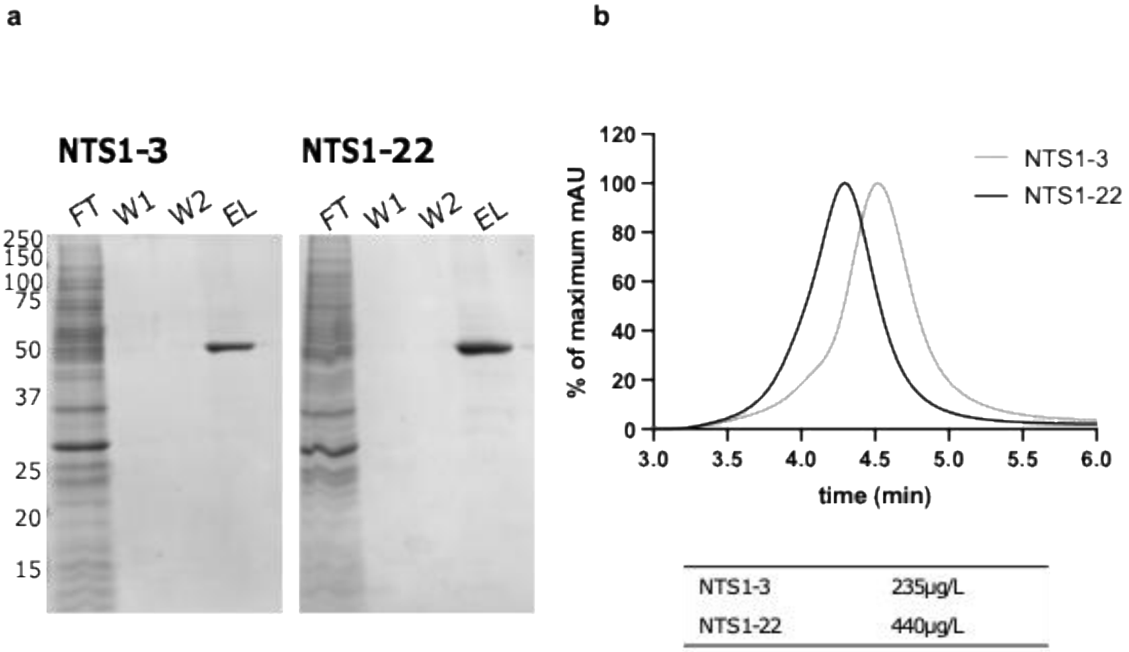


| | |
|---|---|
| NTS1-3 | 235µg/L |
| NTS1-22 | 440µg/L |

**Supplementary Fig. S3. NTS1 purification.** (a) Coomassie-stained 12% polyacrylamide gel of immobilized metal affinity chromatography (IMAC) purified receptor. FT= flow through, W1= wash 1, W2= wash 2, EL= elution 2. NTS1 constructs are identified above. Protein size marker (kDa) is shown on the left. (b) Size exclusion chromatography analysis of the IMAC eluates. The table summarizes the purification yield of NTS1 in the presence of NT(8-13).

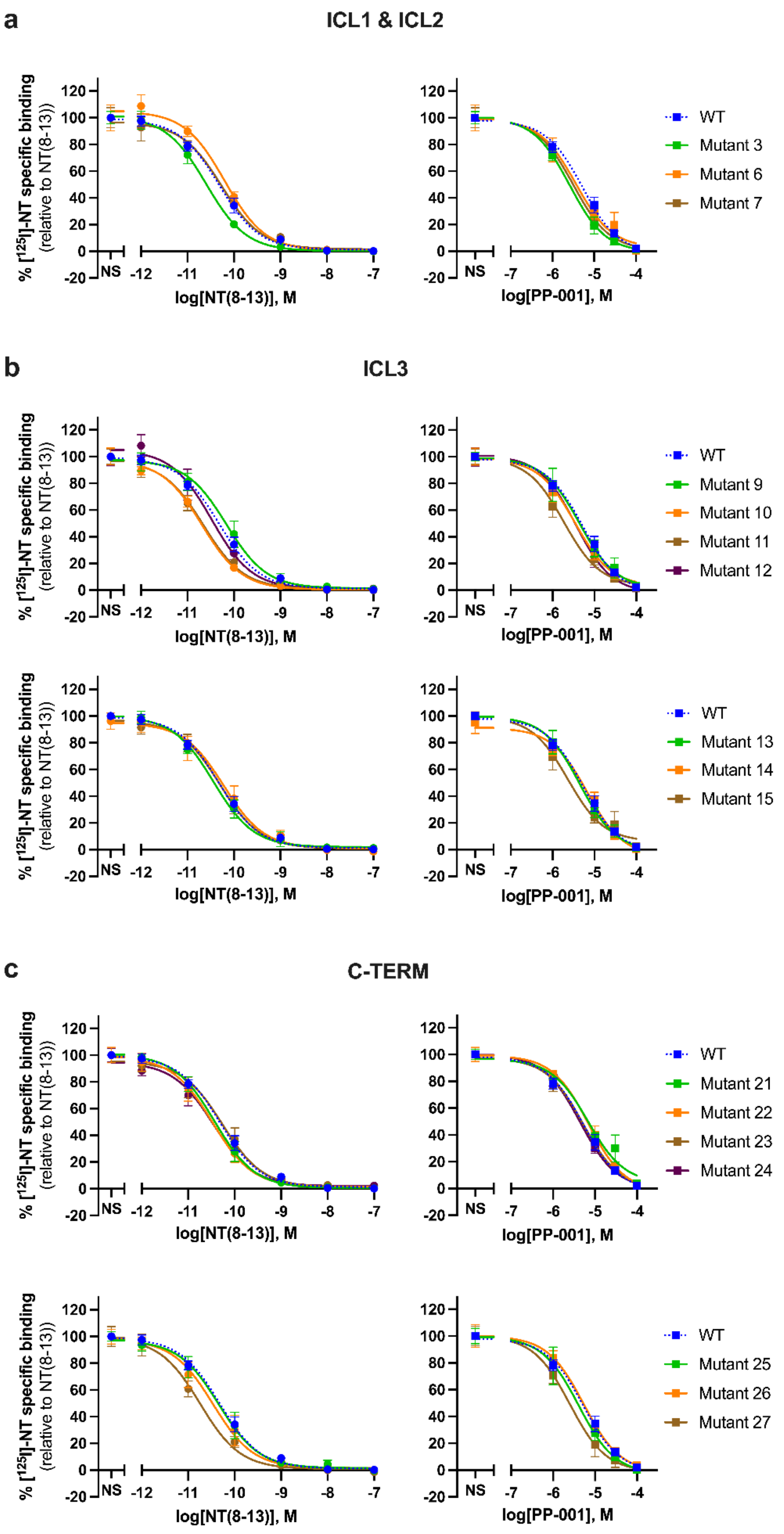
a
ICL1 & ICL2
% [125I]-NT specific binding (relative to NT(8-13))
log[NT(8-13)], M
log[PP-001], M
NS
WT
Mutant 3
Mutant 6
Mutant 7
b
ICL3
Mutant 9
Mutant 10
Mutant 11
Mutant 12
Mutant 13
Mutant 14
Mutant 15
c
C-TERM
Mutant 21
Mutant 22
Mutant 23
Mutant 24
Mutant 25
Mutant 26
Mutant 27

**Supplementary Fig. S4. PP-001-driven inhibition of NT binding is conserved in some NTS1 ICL mutants.** Displacement of $^{125}$I-radiolabeled NT by increasing concentrations of unlabeled NT(8-13) (left) or PP-001 (right), in HEK293 cell membranes transiently transfected with the HA-hNTS1 WT receptor or the NTS1 ICL mutant series (Mutants 1-27). Mutants with substitutions in the ICL1 and ICL2 are presented in (a); ICL3 in (b) and C-terminal in (c). Data represent mean ± SEM of two independent experiments, tested in duplicate.

### Supplementary Table S1. PP-001's effect on NT's potency and efficacy in BRET$^2$ signaling assays

| | $G\alpha_q$ | | $G\alpha_{oA}$ | | $G\alpha_{13}$ | | βarr1 | | βarr2 | |
|---|---|---|---|---|---|---|---|---|---|---|
| **[PP-001], μM** | $EC_{50}$ (nM) | $E_{max}$ (%) | $EC_{50}$ (nM) | $E_{max}$ (%) | $EC_{50}$ (nM) | $E_{max}$ (%) | $EC_{50}$ (nM) | $E_{max}$ (%) | $EC_{50}$ (nM) | $E_{max}$ (%) |
| **0** | 1.4 ± 0.1 | 100 ± 2 | 2.8 ± 0.3 | 100 ± 4 | 1.92 ± 0.07 | 100 ± 1 | 1.91 ± 0.06 | 100 ± 2 | 1.82 ± 0.03 | 100 ± 2 |
| **1** | 1.2 ± 0.1 | 100 ± 2 | 3.0 ± 0.5 | 111 ± 5 | 2.2 ± 0.2 | 100 ± 1 | 1.8 ± 0.1 | 100 ± 3 | 1.7 ± 0.2 | 98 ± 2 |
| **10** | 3.2 ± 0.4 | 97 ± 3 | 3 ± 1 | 109 ± 6 | 4.0 ± 0.5 | 97 ± 3 | 3.9 ± 0.5 | **86 ± 3** | 3.5 ± 0.9 | **75 ± 9** |
| **32** | — | — | 10 ± 10 | 111 ± 5 | 5 ± 7 | 105 ± 5 | **16 ± 13** | **40 ± 20** | 6 ± 3 | **60 ± 20** |
| **100** | **7 ± 9** | 120 ± 10 | 30 ± 40 | 107 ± 12 | **10 ± 20** | 98 ± 3 | **15 ± 15** | **9 ± 7** | 20 ± 20 | **20 ± 20** |

Mean ± SEM. —, not tested. Values in bold are statistically different from NT condition alone ([PP-001] = 0 mM). Kruskal-Wallis test followed by Dunn's correction for multiple comparisons.

**Supplementary Table S2. Effect of negative control compounds on NT's potency and efficacy in $BRET^2$ signaling assays**

| | | $G\alpha_{13}$ | | βarr1 | | βarr2 | |
|---|---|---|---|---|---|---|---|
| | [Compound], μM | $EC_{50}$ (nM) | $E_{max}$ (%) | $EC_{50}$ (nM) | $E_{max}$ (%) | $EC_{50}$ (nM) | $E_{max}$ (%) |
| **NP-001** | 0 | 1.5 ± 0.1 | 100 ± 2 | 1.36 ± 0.08 | 100 ± 2 | 1.31 ± 0.08 | 100 ± 4 |
| | 1 | 1.4 ± 0.1 | 97 ± 1 | 1.44 ± 0.09 | 97 ± 2 | 1.29 ± 0.08 | 97 ± 2 |
| | 10 | 1.4 ± 0.1 | 99 ± 2 | 1.6 ± 0.1 | 97 ± 3 | 1.35 ± 0.09 | 99 ± 2 |
| | 100 | 1.6 ± 0.3 | 100 ± 1 | 1.34 ± 0.09 | 100 ± 3 | 1.4 ± 0.1 | 98 ± 2 |
| **PP-SCR-001** | 0 | 1.0 ± 0.1 | 100 ± 2 | 1.46 ± 0.08 | 100.0 ± 0.9 | 1.4 ± 0.1 | 100.0 ± 0.8 |
| | 1 | 1.6 ± 0.3 | 101 ± 1 | 1.8 ± 0.1 | 103 ± 3 | 1.3 ± 0.1 | 98 ± 3 |
| | 10 | 2.1 ± 0.2 | 101 ± 2 | 1.9 ± 0.2 | 107 ± 5 | 1.8 ± 0.2 | 100 ± 2 |
| | 100 | 3.3 ± 0.4 | 107 ± 1 | 3.4 ± 0.4 | 115 ± 8 | 3.8 ± 0.6 | 93 ± 3 |
| **Palmitate** | 0 | 1.49 ± 0.09 | 100 ± 1 | 1.58 ± 0.07 | 100 ± 3 | 1.55 ± 0.08 | 100 ± 2 |
| | 1 | 2.0 ± 0.1 | 101 ± 2 | 1.8 ± 0.1 | 97 ± 2 | 1.6 ± 0.1 | 98 ± 2 |
| | 10 | 1.9 ± 0.1 | 98 ± 2 | 2.1 ± 0.1 | 99 ± 3 | 1.4 ± 0.1 | 99 ± 3 |
| | 100 | 2.2 ± 0.2 | 93 ± 2 | 1.7 ± 0.1 | 102 ± 2 | 1.6 ± 0.1 | 103 ± 4 |

### Supplementary Table S3. Effect of compounds on homomer and heteromer assay parameters

| | $NTS1_{RlucII}$ / $NTS1_{GFP10}$ | | $NTS1_{RlucII}$ / $APJ_{GFP10}$ | | $APJ_{RlucII}$ / $NTS1_{GFP10}$ | |
|---|---|---|---|---|---|---|
| | $BRET_{50}$ ([GFP10]/[RlucII]) | $BRET_{max}$ ($\Delta BRET^2$) | $BRET_{50}$ ([GFP10]/[RlucII]) | $BRET_{max}$ ($\Delta BRET^2$) | $BRET_{50}$ ([GFP10]/[RlucII]) | $BRET_{max}$ ($\Delta BRET^2$) |
| **Vehicle** | 3.1 ± 0.3 | 0.138 ± 0.007 | 10.3 ± 0.3 | 0.281 ± 0.009 | 4.9 ± 0.2 | 0.125 ± 0.008 |
| **NT(8-13)** | 5.3 ± 0.9 | 0.13 ± 0.01 | 10.1 ± 0.5 | **0.185 ± 0.009** | **5.9 ± 0.5** | 0.109 ± 0.007 |
| **Ape-13** | — | — | 11.6 ± 0.6 | 0.27 ± 0.01 | 4.1 ± 0.4 | 0.13 ± 0.01 |
| **PP-001** | 2.3 ± 0.3 | **0.173 ± 0.007** | 10.0 ± 0.7 | **0.35 ± 0.02** | 4.3 ± 0.3 | **0.16 ± 0.01** |
| **NP-001** | 2.0 ± 0.3 | 0.141 ± 0.008 | 9.8 ± 0.9 | 0.29 ± 0.02 | 3.7 ± 0.3 | 0.112 ± 0.007 |
| **PP-SCR-001** | 2.0 ± 0.3 | 0.14 ± 0.01 | 10.4 ± 0.8 | 0.29 ± 0.02 | 4.8 ± 0.3 | 0.128 ± 0.001 |

Values represent mean ± SEM. —, not tested. Values in bold are statistically different from vehicle condition.

**Supplementary Table S4. Effect of [Ala$^2$]PP-001 and [Ala$^{13}$]PP-001 on NT's potency and efficacy in $BRET^2$ signaling assays**

| | **[Ala$^2$]PP-001** | | | | **[Ala$^{13}$]PP-001** | | | |
|---|---|---|---|---|---|---|---|---|
| | **G$\alpha_{13}$** | | **βarr2** | | **G$\alpha_{13}$** | | **βarr2** | |
| **[Compound], μM** | $EC_{50}$ (nM) | $E_{max}$ (%) | $EC_{50}$ (nM) | $E_{max}$ (%) | $EC_{50}$ (nM) | $E_{max}$ (%) | $EC_{50}$ (nM) | $E_{max}$ (%) |
| **0** | 1.3 ± 0.3 | 100 ± 1 | 2.4 ± 0.1 | 100 ± 3 | 2.5 ± 0.6 | 100 ± 3 | 2.8 ± 0.1 | 100 ± 2 |
| **1** | 1.6 ± 0.4 | 103 ± 2 | 2.9 ± 0.3 | 94 ± 2 | 3.8 ± 0.8 | 103 ± 1 | 3.9 ± 0.2 | 95 ± 2 |
| **10** | 1.5 ± 0.3 | 106 ± 3 | **4.7 ± 0.8** | 102 ± 9 | 10 ± 3 | 112 ± 7 | 21 ± 2 | 83 ± 4 |
| **32** | 1.6 ± 0.4 | 108 ± 3 | **4.3 ± 0.9** | 90 ± 10 | 5 ± 3 | **117 ± 8** | **52 ± 14** | **64 ± 7** |
| **100** | 1.9 ± 0.6 | 110 ± 5 | 5 ± 1 | 100 ± 20 | 15 ± 6 | **116 ± 7** | **150 ± 50** | **72 ± 6** |

Mean ± SEM. Values in bold are statistically different to NT condition alone ([Ala$^x$]PP-001 = 0).

**Supplementary Table S5. Mutations of rNTS1-DARpin22 compared to wild type rNTS1**

| Sequential | B-W | Wild type rNTS1 | rNSTS1-DARpin22 |
|---|---|---|---|
| 83 | 1.51 | S | G |
| 86 | 1.54 | A | L |
| 101 | 2.38 | T | R |
| 103 | 2.40 | H | D |
| 105 | 2.42 | H | Y |
| 119 | 2.56 | L | F |
| 121 | 2.58 | M | L |
| 143 | 3.26 | R | K |
| 161 | 3.44 | A | V |
| 167 | 3.50 | R | L |
| 213 | 4.69 | R | L |
| 234 | 5.35 | R | L |
| 235 | 5.36 | K | R |
| 240 | 5.41 | V | L |
| 253 | 5.54 | I | A |
| 260 | 5.61 | I | A |
| 262 | 5.63 | N | R |
| 263 | 5.64 | K | R |
| 305 | 6.32 | H | R |
| 332 | 6.59 | C | V |
| 342 | 7.26 | F | A |
| 354 | 7.38 | T | S |

**Supplementary Table S6. Mutations introduced in the hNTS1 ICL mutant series**

| | | |
|---|---|---|
| **ICL1** | **A89-H102 (14 AA)** | **ARKKS LQSLQ STVH** |
| Mutant 1 | R90A/K91A/K92A/S93A | AAAAA LQSLQ STVH |
| Mutant 2 | L94A/Q95A/S96A/L97A/Q98A | ARKKS AAAAA STVH |
| Mutant 3 | S99A/T100A/V101A/H102A | ARKKS LQSLQ AAAA |
| **ICL2** | **E165-R184 (20 AA)** | **ERYLA ICHPF KAKTL MSRSR** |
| Mutant 4 | E165A/R166A/Y167A/L168A | AAAAA ICHPF KAKTL MSRSR |
| Mutant 5 | I170A/C171A/H172A/P173A/F174A | ERYLA AAAAA KAKTL MSRSR |
| Mutant 6 | K175A/K177A/T178A/L179A | ERYLA ICHPF AAAAA MSRSR |
| Mutant 7 | M180A/S181A/R182A/S183A/R184A | ERYLA ICHPF KAKTL AAAAA |
| **ICL3** | **A260-R303 (44 AA)** | **ANKLT VMVRQ AAEQG QVCTV GGEHS TFSMA IEPGR VQALR HGVR** |
| Mutant 8 | N261A/K262A/L263A/T264A | AAAAA VMVRQ AAEQG QVCTV GGEHS TFSMA IEPGR VQALR HGVR |
| Mutant 9 | V265A/M266A/V267A/R268A/Q269A | ANKLT AAAAA AAEQG QVCTV GGEHS TFSMA IEPGR VQALR HGVR |
| Mutant 10 | E272A/Q273A/G274A | ANKLT VMVRQ AAAAA QVCTV GGEHS TFSMA IEPGR VQALR HGVR |
| Mutant 11 | Q275A/V276A/C277A/T278A/V279A | ANKLT VMVRQ AAEQG AAAAA GGEHS TFSMA IEPGR VQALR HGVR |
| Mutant 12 | G280A/G281A/E282A/H283A/S284A | ANKLT VMVRQ AAEQG QVCTV AAAAA TFSMA IEPGR VQALR HGVR |
| Mutant 13 | T285A/F286A/S287A/M288A | ANKLT VMVRQ AAEQG QVCTV GGEHS AAAAA IEPGR VQALR HGVR |
| Mutant 14 | I290A/E291A/P292A/G293A/R294A | ANKLT VMVRQ AAEQG QVCTV GGEHS TFSMA AAAAA VQALR HGVR |
| Mutant 15 | V295A/Q296A/L298A/R299A | ANKLT VMVRQ AAEQG QVCTV GGEHS TFSMA IEPGR AAAAA HGVR |
| Mutant 16 | H300A/G301A/V302A/R303A | ANKLT VMVRQ AAEQG QVCTV GGEHS TFSMA IEPGR VQALR AAAA |
| **CTERM** | **N365-Y418 (54 AA)** | **NLVSA NFRHI FLATL ACLCP VWRRR RKRPA FSRKA DSVSS NHTLS SNATR ETLY** |
| Mutant 17 | N365A/L366A/V367A/S368A | AAAAA NFRHI FLATL ACLCP VWRRR RKRPA FSRKA DSVSS NHTLS SNATR ETLY |
| Mutant 18 | N370A/F371A/R372A/H373A/I374A | NLVSA AAAAA FLATL ACLCP VWRRR RKRPA FSRKA DSVSS NHTLS SNATR ETLY |
| Mutant 19 | F375A/L376A/T378A/L379A | NLVSA NFRHI AAAAA ACLCP VWRRR RKRPA FSRKA DSVSS NHTLS SNATR ETLY |

| | | |
|---|---|---|
| Mutant 20 | C381A/L382A/C383A/P384A | NLVSA NFRHI FLATL AAAAA VWRRR RKRPA FSRKA DSVSS NHTLS SNATR ETLY |
| Mutant 21 | V385A/W386A/R387A/R388A/R389A | NLVSA NFRHI FLATL ACLCP AAAAA RKRPA FSRKA DSVSS NHTLS SNATR ETLY |
| Mutant 22 | R390A/K391A/R392A/P393A | NLVSA NFRHI FLATL ACLCP VWRRR AAAAA FSRKA DSVSS NHTLS SNATR ETLY |
| Mutant 23 | F395A/S396A/R397A/K398A | NLVSA NFRHI FLATL ACLCP VWRRR RKRPA AAAAA DSVSS NHTLS SNATR ETLY |
| Mutant 24 | D400A/S401A/V402A/S403A/S404A | NLVSA NFRHI FLATL ACLCP VWRRR RKRPA FSRKA AAAAA NHTLS SNATR ETLY |
| Mutant 25 | N405A/H406A/T407A/L408A/S409A | NLVSA NFRHI FLATL ACLCP VWRRR RKRPA FSRKA DSVSS AAAAA SNATR ETLY |
| Mutant 26 | S410A/N411A/T413A/R414A | NLVSA NFRHI FLATL ACLCP VWRRR RKRPA FSRKA DSVSS NHTLS AAAAA ETLY |
| Mutant 27 | E415A/T416A/L417A/Y418A | NLVSA NFRHI FLATL ACLCP VWRRR RKRPA FSRKA DSVSS NHTLS SNATR AAAA |

**Supplementary Table S7. Effect of ICL mutations on PP-001-driven inhibition of NT binding.**

| | NT(8-13) | PP-001 | |
|---|---|---|---|
| | $IC_{50}$ (pM) | $IC_{50}$ (µM) | Max inhibition (%) |
| WT | 48 ± 3 | 5.3 ± 0.5 | 98 ± 1 |
| Mutant 1 | — | — | — |
| **Mutant 2** | **58 ± 6** | **14 ± 2** | **95 ± 1** |
| Mutant 3 | 25 ± 3 | 2.6 ± 0.4 | 98.6 ± 0.8 |
| Mutant 4 | — | — | — |
| Mutant 5 | — | — | — |
| Mutant 6 | 63 ± 9 | 4 ± 1 | 99.5 ± 0.8 |
| Mutant 7 | 54 ± 8 | 3.1 ± 0.6 | 99 ± 2 |
| Mutant 8 | — | — | — |
| Mutant 9 | 69 ± 8 | 4 ± 1 | 97 ± 2 |
| Mutant 10 | 20 ± 2 | 3 ± 1 | 98 ± 1 |

| | | | |
|---|---|---|---|
| Mutant 11 | 22 ± 3 | 1.9 ± 0.4 | 98.2 ± 0.6 |
| Mutant 12 | 33 ± 6 | 3.6 ± 0.6 | 98.5 ± 0.3 |
| Mutant 13 | 36 ± 4 | 4.4 ± 0.8 | 97.2 ± 0.4 |
| Mutant 14 | 60 ± 9 | 7 ± 1 | 99.4 ± 0.7 |
| Mutant 15 | 50 ± 6 | 2.3 ± 0.7 | 97.6 ± 0.5 |
| **Mutant 16** | **35 ± 3** | **13 ± 2** | **91.9 ± 0.9** |
| **Mutant 17** | **30 ± 2** | **23 ± 3** | **89.2 ± 0.8** |
| Mutant 18 | — | — | — |
| **Mutant 19** | **47 ± 6** | **70 ± 30** | **91 ± 2** |
| **Mutant 20** | **52 ± 6** | **9 ± 3** | **76 ± 6** |
| Mutant 21 | 37 ± 3 | 7 ± 2 | 96.3 ± 0.7 |
| Mutant 22 | 33 ± 4 | 7 ± 1 | 96 ± 1 |
| Mutant 23 | 55 ± 6 | 5.1 ± 0.6 | 96.9 ± 0.9 |
| Mutant 24 | 35 ± 5 | 4.4 ± 0.5 | 97.2 ± 0.7 |
| Mutant 25 | 46 ± 6 | 4 ± 1 | 100 ± 2 |
| Mutant 26 | 34 ± 5 | 5.4 ± 0.9 | 96.6 ± 0.7 |
| Mutant 27 | 34 ± 3 | 2.5 ± 0.5 | 100 ± 1 |

Values represent mean ± SEM. —, unable to generate reliable data.
